# Rice OsPLATZ3 controls ROS homeostasis by inhibiting ROS-Scavenging Activity during Tapetum Degeneration

**DOI:** 10.1101/2024.03.04.583394

**Authors:** Yuanya Li, Jing Wang, Fengxian Tang, Lin Li, Xingyu Cheng, Xialing Sun, Shuangshuang Yu, Pan Xia, Yuxiang Wang, Mingyang Tong, Lizhong Cheng

**Affiliations:** College of Life Science, Yunnan University, Kunming 650091, China

**Keywords:** Rice (*Oryza sativa*), *OsPLATZ3*, tapetum PCD, ROS homeostasis, *OsAPX9*

## Abstract

OsPLATZ3, a transcription factor of rice (*Oryza sativa*) belonging to the PLATZ family, is involved in the repression of transcriptional activity. *PLATZ3* exhibits preferential expression in the tapetal cells during anthers development, regulating tapetum degeneration by controlling the dynamics of reactive oxygen species (ROS) in rice male reproductive processes. The *Osplatz3* exhibits enlarged tapetum with delayed degeneration, significantly lower ROS levels from S5 to S12 in anthers compared to the wild-type, resulting in complete male sterility. In the anthers of *Osplatz3*, there are significant changes in the regulators of programmed cell death (PCD)-induced tapetal degradation, such as *TDR*, *EAT1*, and their effectors *OsCP1*, *OsC6*, *OsAP25*, and *OsAP37*. These findings indicate that *OsPLATZ3* is a new addition to the tapetum PCD regulatory network. In the late stages of anthers development, *Osplatz3* exhibits phenotypes associated with oxidative stress. Transcriptome analysis revealed significant differential expression in 229 genes associated with maintaining ROS homeostasis. Furthermore, Electrophoretic Mobility Shift Assay (EMSA) demonstrate that OsPLATZ3 can bind to the promoter region of *OsAPX9*, a gene encoding a plant ascorbate peroxidase domain protein with putative ROS scavenging capabilities. These findings suggest that OsPLATZ3 serves as a crucial regulatory factor in tapetum PCD by suppressing ROS scavenging activity.

## Introduction

The development of stamens, the male reproductive organs, and the subsequent generation of pollen grains are critical components of male reproductive growth in flowering plants, involving complex physiological and biochemical processes (Mondol et al., 2019). The anther of rice consists of four distinct layers: the epidermis (outermost layer), endothecium, middle layer, and tapetum (innermost layer) (Bai et al., 2019). The tapetum cells, located within the sporophytic layer, play a crucial role in the process of pollen formation. These cells are responsible for breaking down callose, allowing the release of microspores from tetrads (Wu and Cheung, 2000). Additionally, tapetum cells support the growth of microspores by providing essential nutrients and serving as reservoirs for sporopollenin precursors required for sexine production (Bai et al., 2019).

The precise timing of tapetal cell death, which occurs through developmentally regulated programmed cell death (PCD), is critical for pollen development (Xie et al., 2014). Tapetal PCD is intricately regulated by evolutionarily conserved transcriptional cascades. Several transcription factors have been identified in rice as being involved in tapetum PCD, including *Undeveloped Tapetum 1* (*UDT1*) (Jung et al., 2005), *TAPETUM DEGENERATION RETARDATION* (*TDR*) (Li et al., 2006; Ji et al., 2013; Niu et al., 2013; Fu et al., 2014; Ko et al., 2014; Chen et al., 2017), *TDR INTERACTING PROTEIN2 (TIP2)* (Fu et al., 2014; Ko et al., 2014), *ETERNAL TAPETUM 1* (*EAT1*) (Ji et al., 2013; Niu et al., 2013), *GAMYB* (Aya et al., 2009), *PERSISTANT TAPETAL CELL1* (*PTC1*) (Li et al., 2011), *OsMADS3* (Hu et al., 2011), as well as their downstream *DEFECTIVE TAPETUM CELL DEATH 1* (*DTC1*) (Yi et al., 2016). Downstream of OsGAMYB, TDR and PTC1 activate the expression of lipid transfer protein OsC6, thus promoting pollen wall formation and tapetal cell PCD (Li et al., 2006; Zhang et al., 2010; Li et al., 2011). UDT1 (Jung et al., 2005) is a homolog of bHLH142 (Ko et al., 2014), and bHLH142 promotes PCD through *TDR* and EAT1 (Niu et al., 2013). TIP2 interacts with TDR and sequentially regulates the expression of EAT1 (Fu et al., 2014; Ko et al., 2014). EAT1 is essential for the synthesis of *OsAP25* (an aspartic protease in rice) and *OsAP37* during the tapetum PCD (Niu et al., 2013). TDR interacts with EAT1, and TDR is necessary to initiate *Cysteine Protease 1* (*OsCP1*) (Li et al., 2006). The proteins OsCP1, OsAP25, and OsAP37 play critical roles in the induction of tapetal PCD in rice (Lee et al., 2004; Niu et al., 2013).

Reactive Oxygen Species (ROS) play a vital role as a key regulator in initiating PCD during various biological processes such as tapetum degeneration, aleurone degradation, aerenchyma tissue formation, seed growth and germination, and tracheary element development (Yi et al., 2016). ROS levels that are too low or too high can hinder plant growth and development (Mittler, 2017). Therefore, it is essential to maintain a balance between the generation and elimination of intracellular ROS. Plants manage a dynamic equilibrium of ROS through enzymatic or non-enzymatic pathways and have developed a comprehensive antioxidant system (Maruta et al., 2016). In the tapetal PCD regulatory system, some key genes control the tapetum degeneration by regulating the ROS dynamics during the male reproductive process of rice. For example, DTC1 regulates ROS levels by binding to metallothionein during tapetum degeneration (Yi et al., 2016). Additionally, MADS3 interacts with MT-1-4b to regulate ROS homeostasis throughout the later stages of anther development. Transgenic plants with reduced MT-1-4b expression have lower pollen fertility and increased amounts of superoxide anion (Hu et al., 2011).

PLATZs are plant-specific DNA-binding proteins that contain zinc-binding domains and are involved in inhibiting transcription. These proteins identify and interact with A/T-rich DNA sequences to carry out their regulatory function (Nagano, 2001). In Arabidopsis, RITF1, a member of the PLANT RICH SEQUENCE and ZINC-BINDING TRANSCRIPTION FACTOR (PLATZ) family, plays a crucial role in regulating root development by modulating ROS levels (Yamada et al., 2019). AtPLATZ2 interacts with and inhibits the activity of calcium-binding proteins involved in salt stress signaling pathways (Liu et al., 2020). *ORESARA15* (*ORE15*) regulates the size of the root apical meristem (RAM) by balancing the actions of auxin and cytokinin signaling pathways (Timilsina et al., 2022). *PLATZ4* enhances drought resistance by promoting stomatal closure and regulating genes associated with abscisic acid (ABA) signaling (Liu et al., 2023). In rice, the genes *GL6* and *SG6* belong to the PLATZ protein family and are involved in controlling the length and size of rice grain (Wang et al., 2019). In Maize (*Zea mays*), the *fl3* gene encoding a PLATZ protein is a semidominant negative mutant affecting the growth of endosperm and storage reserves in seeds (Li et al., 2017). *ZmPLATZ2,* a gene found in maize, is associated with starch production and specifically attaches to the promoter region of the *ZmSSI* gene (Li et al., 2021). In wheat, *RHT25,* also known as PLATZ-A1, plays a crucial role in controlling the sensitivity to the plant hormone gibberellin (GA) and influencing plant height during stem elongation and spike development stages (Zhang et al., 2023). Despite the critical importance of the PLATZ family in various physiological functions, little is known about the molecular mechanisms by which the PLATZ family regulates ROS homeostasis in rice, which activates the tapetum PCD.

In this study, we found that *OsPLATZ3* plays a crucial role in regulating rice tapetum PCD by modulating the ROS homeostasis. *OsPLATZ3* is expressed in the tapetal cells of developing anthers. The *Osplatz3* exhibits delayed tapetum degeneration due to reduced levels of ROS in the late stage of anthers development, resulting in male sterility. The abnormal decrease of ROS levels in *Osplatz3* may be attributed to altered gene expression involved in maintaining ROS homeostasis. Furthermore, we demonstrated that OsPLATZ3 can bind to the promoter region of *OsAPX9* (*Os04g51300*). *OsAPX9* is a rice ascorbate peroxidase domain-containing protein, which is homologous to Arabidopsis ROS scavenging gene *APX4*. These findings provide insights into the function of *OsPLATZ3* in the regulation of tapetum PCD in the rice, suggesting that OsPLATZ3 controls ROS levels partly through *OsAPX9*.

## Results

### Identification of *Osplatz3* mutant resulting in absolute male sterility

We obtained a male sterile rice mutant, strain 1C-04607, from the T-DNA mutant pool (Jeon et al., 2000) and named it *Osplatz3-1*. The T-DNA was inserted at 26 bp of the second exon of *Os02g07650*, resulting in a homozygous male-sterile phenotype in *Osplatz3-1* (Fig. 1, A-D, I and L). Furthermore, we identified another T-DNA strain 4A-02615 of the *Os02g07650* gene, named *Osplatz3-2*, with the T-DNA inserted at 66 bp of the second exon, leading to a heterozygous fertile phenotype (Fig. 1, A, E-G, J and M). When the *Osplatz3-2* heterozygous progeny were selfed, the normal to sterile plant ratio was approximately 3:1 (67 fertile:29 sterile plants: x^2^=0.52; P>0.05), suggesting that the sterile phenotype is governed by a single recessive mutation. All sterile progeny exhibited homozygous for the T-DNA insertion, approximately one-third of the fertile progeny were wild-type and lacked T-DNA, whereas the remaining carried T-DNA on only one chromosome. This indicates that the observed phenotype results from the insertion of T-DNA.

**Figure 1.**
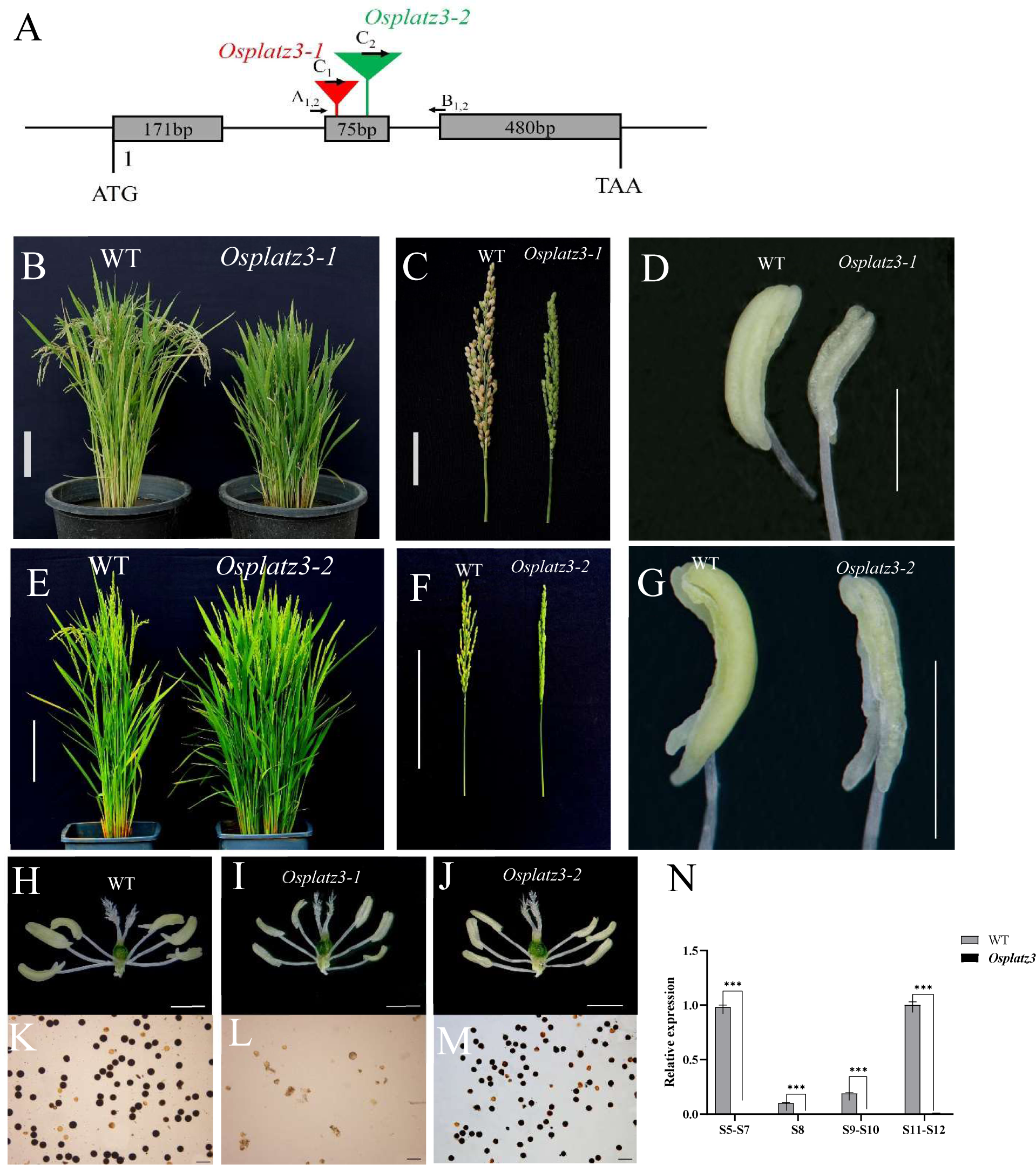
Characterization of T-DNA insertion mutants in *OsPLATZ3*. A, Gene structure of OsPLATZ3. Gray boxes represent exons, lines connecting boxes represent introns and UTRs, red and green triangles indicate T-DNA insertions of *Osplatz3-1* (line 1C-04607) (red) and *Osplatz3-2* (line 4A-02615) (green). Arrows A_1,2_, B_1,2_, and C_1,2_ show the left, right primers and vector border primers for RT-PCR analysis used for identification of *Osplatz3-1* and *Osplatz3-2* mutants. B, Wild-type and *Osplatz3-1* (line 1C-04607) mature plants. C, Wild-type and *Osplatz3-1* spikes. D, Wild-type and *Osplatz3-1* anthers. E, Wild-type and *Osplatz3-2* mature plants. F, Wild-type and *Osplatz3-2* spikes. G, Wild-type and *Osplatz3-2* anthers. H, Wild-type stamens. I, *Osplatz3-1* stamens. J, *Ospla tz3-2* stamens. K, mature pollen grains of wild-type. L, pollen remnants of of *Osplatz3-1*. M, mature pollen grains of *Osplatz3-2*. N. qRT-PCR analysis of OsPLATZ3 expression in different developmental stages of *Osplatz3* inflorescence. Bars = 15 cm in B, E, and F. Bars = 5 cm in C. Bars = 1mm in D, G, H, I, and J. Bars = 100 μm in K and L. Bars = 50 μm in M.

The growth pattern of *Osplatz3-1* and *Osplatz3-2* is similar to that of wild-type plants during the early stages of vegetative development. However, as *Osplatz3-1* grew and developed, it exhibited plant dwarfism, with a height approximately 80% of the wild-type (Fig.1B). The tillering of *Osplatz3-1* is twice that of the wild-type. Unlike the wild-type, the shriveled spikes of *Osplatz3-1* remained green and erect, with a length of around 90% of the wild-type full spikes (Fig. 1C). The *Osplatz3-1* anthers were small, pale in color, and had a curled, wrinkled, and sponge-like appearance (Fig. 1D). Pollen staining showed that all wild-type pollen grains turned black, indicating nearly full maturation (Fig. 1K), while the *Osplatz3-1* exhibited a completely sterile phenotype, as it lacked pollen grains and only showed remnants of pollen abortion (Fig. 1L). In contrast, the tillering of *Osplatz3-2* is also twice that of the wild-type, but its plant height, panicles, and anthers are basically the same as the wild-type, with only a slightly delayed growth period. *Osplatz3-2* has many pollen grains and is fertile, just like the wild-type (Fig. 1, E-G, J and M).

According to the annotation from the Rice Genome Database, *Os02g07650* codes for a zinc-binding protein belonging to the rice PLATZs family of transcription factors, referred to as *OsPLATZ3* (*Os02g07650*) in our study. During the S5-S12 of anthers development, *OsPLATZ3* showed significantly high expression in the wild-type, as indicated by qRT-PCR analysis (Fig. 1N, Supplemental Table S1 for primers). Compared to the wild-type, the expression levels of *OsPLATZ3* were barely detectable in the *Osplatz3* homozygote at different anther stages (Fig. 1N), suggesting that *OsPLATZ3* has undergone mutation in *Osplatz3*.

### *OsPLATZ3* is mainly expression in the tapetum during the anther development

To comprehend the expression of *OsPLATZ3*, we conducted qRT-PCR analysis. It was observed that *OsPLATZ3* exhibited high expression levels in 10-day roots, 10-day shoot, and all stages of inflorescences, reaching its peak during the S5-S7 (Supplemental Fig. S1A). Following previous studies (Zhang and Wilson, 2009), we delineated stages of rice anther development and selected S5-S7, S8, S9-S10, S11, and S12 for GUS staining (Supplemental Fig. S1B). The GUS staining results showed that from S5 to S12 (Supplemental Fig. S1B), except for the slightly weaker GUS signal in the anthers at S11, there was a strong GUS staining at all other stages, indicating that OsPLATZ3 has a high expression level in various stages of anther development. In addition, GUS staining of the vegetative tissues in the transgenic plants showed that, apart from the GUS signal in the meristematic root tips of adventitious roots (Supplementary Fig. S1C), GUS staining in other mature tissues was either very weak or not visible. This result is consistent with the qRT-PCR analysis (Supplemental Fig. S1A), indicating that OsPLATZ3 is mainly expressed in young tissues such as young roots, seedlings, and various stages of anther development.

During our investigation into *OsPLATZ3* expression in anthers, we conducted RNA in situ hybridization assays. Initially, *OsPLATZ3* transcripts were observed at S7 (Fig. 2A), with their abundance steadily increasing in the tapetal cells and microspore mother cell at S8 (Fig. 2, B and C). From S8 to S11, the abundance of *OsPLATZ3* transcripts in tapetal cells remained consistently high (Fig. 2, B-F). *OsPLATZ3* expression became undetectable at S12 (Fig. 2G). In contrast, the sense probe in the control only detected background signals at S7 (Fig. 2H), which signifies the initial appearance of *OsPLATZ3* transcripts, and at S11 (Fig. 2I), which signifies their final appearance. These results indicate that *OsPLATZ3* is highly expressed during anthers development, primarily in the tapetal cells, and to a lesser extent in microspores.

**Figure 2.**
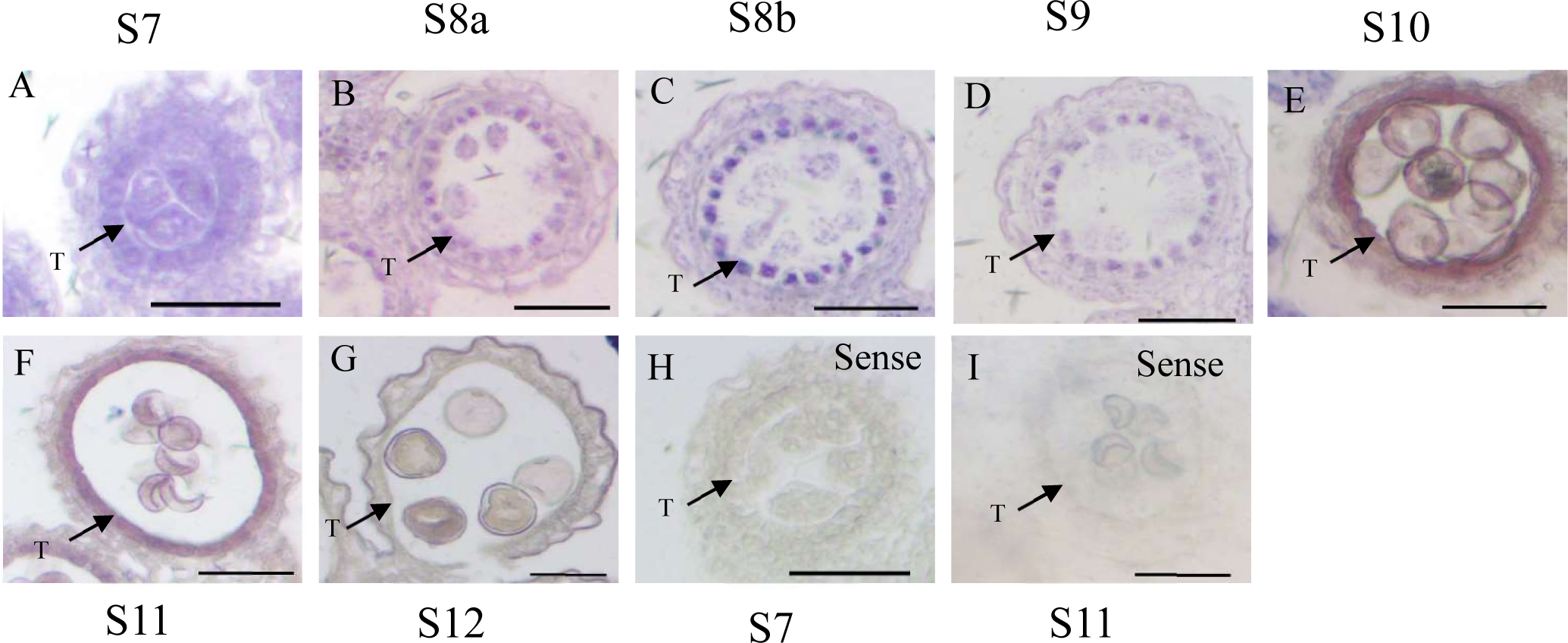
In situ hybridization of *OsPLATZ3* expression in wild-type anthers at different developmental stages. In situ hybridization of OsPLATZ3 transcripts in anthers at S7 (A), S8a (B), S8b (C), S9 (D), S10 (E), S11 (F), and S12 (G). Anthers at S7 (H) and S11 (I) hybridized with the *OsPLATZ3* sense probe. T, tapetum; Bars = 100 μm.

### *OsPLATZ3* mutation cause microspore sterility

To further validate the impact of the *OsPLATZ3* mutation on the anther development, we constructed the *Cas9*-*OsPLATZ3* vector and transformed rice (the edited sites detailed in Supplemental Table S1). qRT-PCR analysis of the homozygous mutants indicated that, compared to the wild-type, the expression of *OsPLATZ3* in the inflorescences during S5-S12 of *Cas9-OsPLATZ3* was extremely low and almost undetectable, demonstrating the successful editing of *OsPLATZ3* (Supplemental Fig. S2).

Subsequently, we examined anther sections from wild-type, *Osplatz3*, and *Cas9-OsPLATZ3*. Based on the previously established classification of anthers development, we categorized the rice anther development process into 12 stages (Zhang and Wilson, 2009). The first mitotic division of the sporogenous cells (Supplemental Fig. S3, A, D, and G) resulted in the production of two secondary sporogenous cells (Supplemental Fig. S3, B, E, and H), which further divided to produce four microspore mother cells (Supplemental Fig. S3, C, F, and I). At this stage, the formation of microspore mother cells (MMC) in *Osplatz3* and *Cas9-OsPLATZ3* were normal and consistent with the wild-type (Supplemental Fig. S3).

At the MMC stage, no significant differences in anther structure were observed between the wild-type, *Osplatz3*, and *Cas9-OsPLATZ3*, with normal epidermis, endothecium, middle layer, tapetum, and microspore mother cells (Fig. 3A, a, h and o). At S8, MMCs of the wild-type underwent meiosis to generate tetrads of four haploid microspores. The tapetal cells experienced substantial changes due to developmental differentiation, and the cytoplasm became particularly concentrated in the middle layer (Fig. 3A, b and c). Meiosis of *Osplatz3* and *Cas9-OsPLATZ3* is normal, similar to the wild-type (Fig. 3A, i, j, p and q). In wild-type anthers, the callose wall was completely degraded by S9, releasing the microspores from the tetrads. The tapetal cell cytoplasm was heavily pigmented but without large vesicles, the middle layer was nearly invisible, and the wild-type microspores were spherical and full (Fig. 3A, d). At S9, the *Osplatz3* and *Cas9-OsPLATZ3* anthers showed substantial differences from the wild-type; the tapetal cell cytoplasm was not deeply stained, and the cells had huge vesicles that began to enlarge dramatically and partly crowding out the space for the microspores (Fig. 3A, k and r). In S10, the tapetum of wild-type degenerates and the uninucleate microspores become more vacuolated and assume a rounded shape, indicating the onset of pollen grain enlargement (Fig. 3A, e). At this stage, compared to the wild-type, *Osplatz3* and *Cas9-OsPLATZ3* show more pronounced phenotypic differences, with the tapetum not undergoing degeneration and exhibiting a darker staining, resembling the wild-type tapetum at the S9, demonstrating a delayed degeneration phenotype. Additionally, although the microspores enlarge, their morphology is irregular (Fig. 3A, l and s). By S11, the wild-type tapetal cells had almost entirely disappeared, and vacuolated pollen transformed into binucleate pollen, accumulating nutrients like starch and liposomes. Starch accumulated internally in binucleate pollen, causing the vesicles to gradually diminish (Fig. 3A, f). At this stage, *Osplatz3* and *Cas9-OsPLATZ3* tapetal cells had not entirely degenerated, and binucleate microspores could no longer form (Fig. 3A, m and t). At S12, the generative cell in the microspore divided into two sperm cells, and the mature pollen underwent two mitotic divisions to become trinucleated. The vesicles vanished completely, and a significant number of starch granules formed in the mature pollen grain (Fig. 3A, g). At this point, *Osplatz3* microspores had been completely degraded, with remnants in the core of the anther locules (Fig. 3A, n), and the associated developmental processes could not be observed for *Cas9-OsPLATZ3*. It is evident from the data that *Osplatz3* and *Cas9-OsPLATZ3* displayed expansion of the tapetum during the stage of uninucleate pollen, thereby encroaching on the space available for microspores. The tapetal cells persisted during the trinucleate pollen stage, resulting in the abortion of microspore development and the visible remnants of degenerated pollen. This signifies a delayed tapetum PCD in *Osplatz3* and *Cas9-OsPLATZ3*, leading to male sterility.

**Figure 3.**
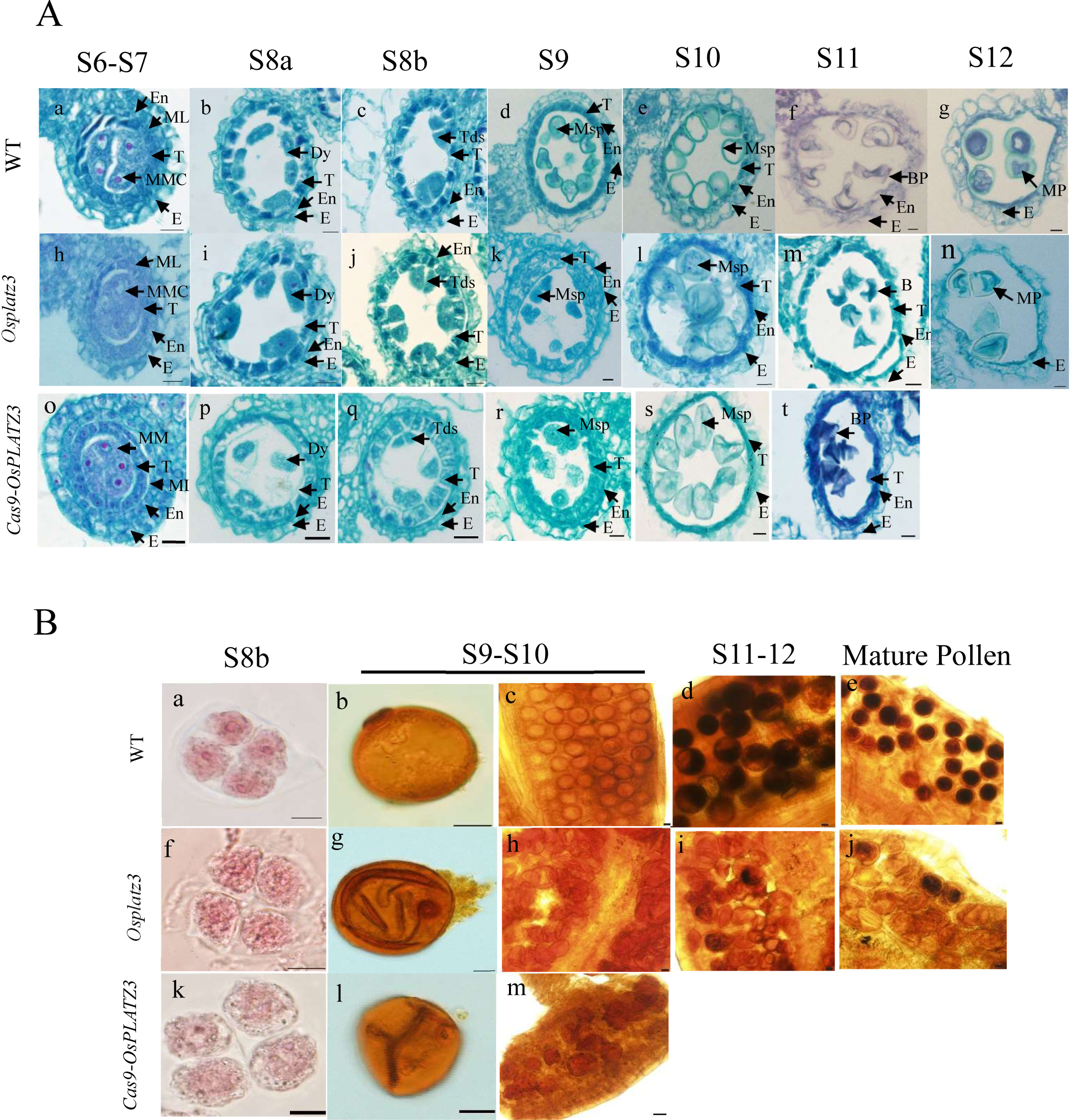
OsPLATZ3 mutation cause microspore sterility. A, Comparison of Anther Development between the Wild-type, *Osplatz3* and *Cas9-OsPLATZ3*. BP, biceullar pollen; Dy, dyad cell; E, epidermis; En, endothecium; MMC, microspore mother cell; ML, middle layer; MP, mature pollen; Msp, microspore parietal cell; T, tapetum; Tds, tetrads. Bars = 10 μm. a, In the S6-S7 of wild-type, the remaining archesporial cells continue to divide, producing four microspore mother cells. b, The first meiotic division (S8a) of wild-type, showing elliptical dyads. c, The second meiotic division (S8b) of wild-type, showing tetrads about to separate. d, In the S9 of wild type, microspores separate from tetrads. e, In the S10 of wild type, microspores show vacuolation and significant increase in volume. f, In the S11 of wild type, uninucleate microspores undergo mitosis, producing binucleate microspores. At this stage, the microspores have a “crescent” shape, and the tapetum completely degenerates. g, In the S12 of wild type, microspores accumulate nutrients and become non-transparent. h and o, In the S6-S7 of *Osplatz3* and *Cas9-OsPLATZ3*, the remaining archesporial cells continue to divide, producing four microspore mother cells. i and p, The first meiotic division (S8a) of *Osplatz3* and *Cas9-OsPLATZ3*, showing elliptical dyads. j and q, The second meiotic division (S8b) of *Osplatz3* and *Cas9-OsPLATZ3*, showing tetrads about to separate. k and r, In the S9 of *Osplatz3* and *Cas9-OsPLATZ3*, microspores separate from tetrads and the tapetum enlarges. l and s, In the S10 of *Osplatz3* and *Cas9-OsPLATZ3*, microspores show vacuolation and significant increase in volume. The roundness of microspores differs from WT. m and t, In the S11 of *Osplatz3* and *Cas9-OsPLATZ3*, uninucleate microspores undergo mitosis, producing binucleate microspores. At this stage the microspores have a “crescent” shape, and the mutant tapetum does not degenerate. n, In the S12 of *Osplatz3*, the shape of the microspores is abnormal, showing a transparent state. It is difficult to find images of the S12 of *Osplatz3*. B, Observation of pollen staining in the wild-type and *Osplatz3*, *Cas9-OsPLATZ3*. a, Tetrad stage of the wild-type, which are about to separate from each other. b, Pollen development in the wild-type at S10. c, Local pollen of the wild-type at S10, with rounded spore surface. d, Local pollen of the wild-type at stages 11-12, spores have nutrient deposits. e, Mature pollens of the wild-type. f and k, Tetrad stage of *Osplatz3*, *Cas9-OsPLATZ3*, which are about to separate from each other. g and l, Pollen development in the S10 of *Osplatz3*, *Cas9-OsPLATZ3*, spores surface is wrinkled. h and m, Local pollen at S10 of *Osplatz3*, *Cas9-OsPLATZ3*, surface of mutant spores is wrinkled. i, Local pollen at S11-S12 of *Osplatz3*, very few spores have nutrient deposits. J, Mature pollens of *Osplatz3*.

In addition, a comparison of pollen viability between *Osplatz3*, *Cas9-OsPLATZ3*, and wild-type confirmed the above findings. Anthers were stained with acetic fuchsin and potassium iodide, revealing that at S8b, WT, *Osplatz3*, and *Cas9-OsPLATZ3* formed typical tetrads (Fig. 3B, a, f and k). However, in S9-S10, unlike the wild-type (Fig. 3B, b and c), irregularly shaped microspores were observed in the anthers of *Osplatz3* and *Cas9-OsPLATZ3* (Fig. 3B, g and l), with nearly all microspores displaying irregular morphology (Fig. 3B, h and m). By S11-S12, *Osplatz3* displayed a lack of accumulation of starch or other nutrients (Fig. 3B, i). Meanwhile, it is possible that *Cas9-OsPLATZ3* microspores had ceased to form. During mature pollen development, *Osplatz3* microspores may have ceased to develop, rendering the microspores sterile (Fig. 3B, j). The above results revealed that the microspores of both *Osplatz3* and *Cas9-OsPLATZ3* were sterile, indicating that the *OsPLATZ3* mutation led to microspore sterility. Furthermore, the examination of the sectioned *Osplatz3* embryos demonstrated normal development of the embryo *sac*, providing additional support for this conclusion (Supplemental Fig. S4).

### OsPLATZ3 is a nucleus-localized transcriptional repressor

To understand the subcellular localization of OsPLATZ3, GFP Nucleus marker was used as a reference, and the pCAMBIA1300-35S-AtHY5-mCherry and OsPLATZ3-GFP vectors were constructed (see Supplemental Table S1 for primers). Confocal laser scanning microscopy showed that the fluorescence signal of the GFP control was distributed in the cell nucleus. Consistently, the yellow fluorescence signal of the OsPLATZ3-GFP fusion protein was also distributed in the cell nucleus. Additionally, fluorescence signals were observed on the cell membrane, suggesting a possible localization on the cell membrane. However, we did not use a marker for cell membrane localization as a control, so we cannot confirm its localization on the cell membrane at this time. Therefore, we believe that OsPLATZ3 is a transcription factor localized in the cell nucleus (supplemental Fig. S5A).

The yeast system was used to verify the transcriptional activation of OsPLATZ3. In the selective SD medium lacking tryptophan and histidine (SD/-Trp-His), only the positive control transformants were able to grow. The pBD-PLATZ3-FL and the pBD (negative control) transformants failed to grow on the SD/-Trp/-His medium (Supplementary Fig. S5B). These results suggest that OsPLATZ3 may have a transcriptional inhibitory effect, which is consistent with previous studies (Nagano, 2001).

Effector vectors containing fusion transcription factors GAL4/*OsPLATZ3*, GAL4/*OsPLATZ3N*, GAL4/*OsPLATZ3C*, and GAL4/VP16-*OsPLATZ3*, along with reporter vectors containing an upstream activation sequence (5ⅩUAS) promoter driving the firefly luciferase (LUC) reporter gene in the GAL4/VP16-5XUAS system, were constructed for evaluating the transcriptional activation activity of *OsPLATZ3*. The results indicated that *OsPLATZ3* acts as a transcriptional inhibitory factor, exhibiting significantly downregulating the activity of the LUC reporter gene when compared to the GAL4 and GAL4/VP16 control (Supplemental Fig. S5C).

### *OsPLATZ3* mutation leads to defective tapetum degradation and reduced ROS accumulation

To investigate the abnormal degradation in *Osplatz3*, we performed terminal deoxynucleotidyl transferase-mediated dUTP nick-end labeling (TUNEL) assay to evaluate the extent of PCD in *Osplatz3*. In wild-type, green fluorescence signals initially emerged at S8a, increased at S8b and S9, weakened at S10, and disappeared completely at S11 (Fig. 4A, a-e). However, in the tapetum of *Osplatz3*, sustained apoptotic signals were observed through green fluorescence from S8 to S11 (Fig. 4A, f-j). Among these, the fluorescent signals were most pronounced during S8 (Fig. 4A, f and g), S10 (Fig.4A, i), and S11 (Fig.4A, j), while slightly weaker during S9 (Fig. 4A, h). This indicates that *OsPLATZ3* mutation disrupts the orderly progression of tapetum PCD, leading to delayed occurrence.

**Figure 4.**
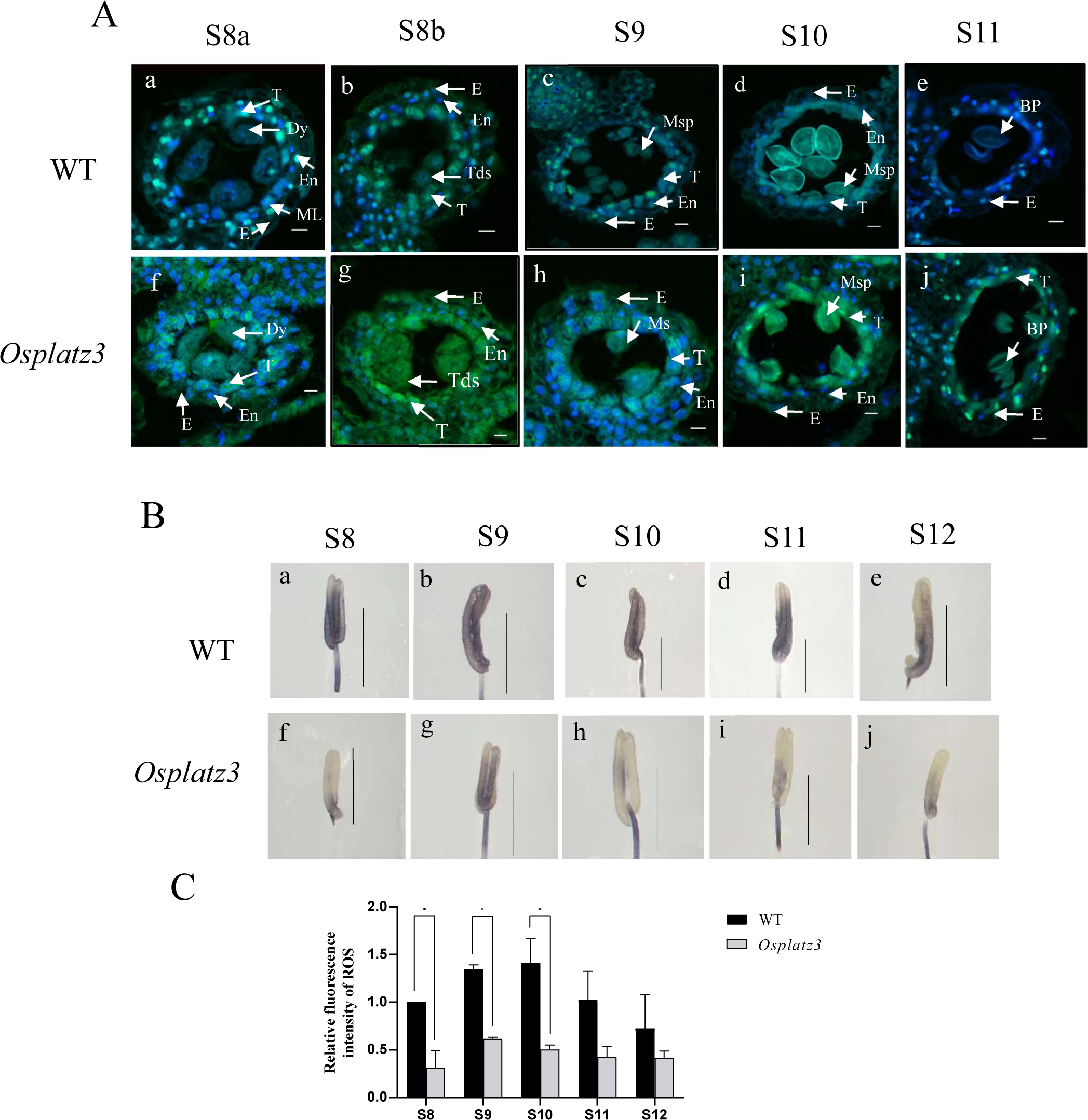
TUNEL assay and analysis of superoxide anion levels. A, TUNEL assay to detect DNA fragmentation. WT (a-e) and *Osplatz3* (f-j) signal in the anthers from S8 to S11. The green fluorescence indicates TUNEL-positive areas. Cell nuclei are stained with DAPI, showing blue fluorescence. BP, bicellular pollen; Dy, dyad cell; E, epidermis; En, endothecium; ML, middle layer; Msp, microspore parietal cell; T, tapetum; Tds, tetrads. Bars = 10 μm. B, NBT staining of superoxide anions from wild-type and *Osplatz3* anthers (from S8 to S12). Bars = 1 mm. C, Distribution of ROS in rice anthers (from S8 to S12) detected by DCFH-DA staining. Relative intensity of ROS fluorescence was calculated using ImageJ software.

To investigate whether *OsPLATZ3* plays a role in ROS homeostasis, we conducted measurements of superoxide anions in wild-type and *Osplatz3* anthers using Nitroblue Tetrazolium (NBT). In wild-type anthers, S9 and S10 exhibited the most extensive staining (Fig. 4B, b-c), while S8, S11, and S12 showed lighter staining (Fig. 4B, a, d and e). In contrast, the *Osplatz3* mutant displayed significantly lower levels of staining compared to the wild-type across all S8-S12 (Fig. 4B, f-j), with only some staining observed at S9 (Fig. 4B, g), indicating reduced ROS levels in the *OsPLATZ3* mutant.

Furthermore, a fluorescence test was conducted using the 2,7-dichlorodihydrofluorescein diacetate (DCFH-DA) assay to quantify total ROS levels in rice anthers (Zhao et al., 2018). ROS levels in *Osplatz3* anthers were notably lower than in the wild-type from S8 to S12 (Fig. 4C). These results suggest the crucial role of optimal ROS levels in PCD, as lower ROS levels in *Osplatz3* anthers led to delayed tapetum degradation.

### *OsPLATZ3* modulates tapetum PCD during late anther development

To gain a deeper understanding of the regulatory functions of *OsPLATZ3*, we carried out RNA sequencing (RNA-seq) to compare genome-wide mRNA levels in wild-type and *Osplatz3* anthers from S5 to S12. We conducted three biological replicates for both wild-type and *Osplatz3*. The anticipated functions of genes were derived from release 7.0 of the Rice Genome Annotation Project (RGAP). The genes that express changes in the *Osplatz3* can be functionally classified into three categories (Supplemental Fig. S6A): Cellular components (1148 genes down-regulated and 863 genes up-regulated), Biological processes (526 genes down-regulated and 469 genes up-regulated), and Molecular functions (344 genes down-regulated and 248 genes up-regulated) (Supplemental Fig.S6B). In S5-S7, S8, S9-S10, S11, and S12, there were 385, 304, 1725, 1843, and 960 genes, respectively, showing significant differential expression (Supplemental Fig. S6C).

According to the identification of 140 specific genes in anthers (Suwabe et al., 2008), we observed significant changes in the expression of 112 genes related to anthers in *Osplatz3* (Supplemental Fig. S7). These findings highlight the crucial role of *OsPLATZ3* in the process of anthers development. *OsPLATZ3* did not have an impact on the expression of specific genes involved in the early development of rice anthers. In contrast, expression levels of genes associated with late anther development showed significant changes. For example, *OSMST8* (*Os01g38670*), responsible for sugar transport, exhibited a 4-fold reduction at S9-S10 and a 7-fold decrease at S11 (Supplemental Table S2). Similarly, *OSINV4* (*Os04g33720*), involved in pollen maturation and starch synthesis, decreased by 5-fold at S9-10, 2-fold at S11, and 13-fold at S12 (Supplemental Table S2). Reduced expression of *OSMST8* and *OSINV4* has been linked to male sterility in rice (Oliver et al., 2007). Furthermore, the expression of four other male sterility-related genes, including *Os04g28620*, *Os04g28520*, *Os09g39410*, and *DPW* (*Os03g07140*), also showed significant variations, supporting the view that *OsPLATZ3* plays a critical role in late anther formation (Supplemental Table S2).

In the transcriptomic data, we observed significant variability in the expression of genes associated with tapetum PCD. For instance, *OsTDR* (Li et al., 2006), *OsEAT1* (Niu et al., 2013), *OsCP1*, *AP25*, *OsAP37* (Lee et al., 2004; Li et al., 2006; Niu et al., 2013), and *OsC6* (Zhang et al., 2010) exhibited notable changes. The expression of *TDR* was observed to be 3 times lower in the *Osplatz3* at the S9-S10 and 3 times lower at the S12 (Supplemental Table S2), both crucial for tapetal development and degeneration. Mutants displayed delayed tapetum PCD (Li et al., 2006). Additionally, the expression of *OsC6* in S9-S10 was 22 times lower, 10 times higher in S11, and 3 times lower at S12 in the *Osplatz3* (Supplemental Table S2). *OsC6* encodes a small mobile lipid transfer protein essential for rice meiotic anther formation (Zhang et al., 2010). *OsCP1* was down-regulated by 4 times at S9-S10 (Supplemental Table S2). These results indicated that *OsPLATZ3* regulates tapetum PCD during late anther formation in rice.

The above results were verified using qRT-PCR (Fig. 5, A-F). We observed a significant decrease in the expression of *TDR* in the *Osplatz3* (Fig.5A). The levels of *EAT1* showed significant variations in the *Osplatz3* (Fig. 5B), suggesting that OsPLATZ3 may be located upstream of *TDR*, *EAT1/DTD*, exerting regulatory effects on *TDR*, *EAT1/DTD*. Furthermore, there were significant differences in the expression levels of *TDR* downstream genes *OsCP1* and *OsC6* at various stages of anther development (Fig. 5, C-D). Additionally, the transcription levels of two aspartic protease-encoding genes, *AP25* and *AP37*, also decreased in *Osplatz3* (Fig. 5, E-F). This suggests that *OsPLATZ3* may affect the genes encoding the effectors of tapetal PCD or secreted proteins.

**Figure 5.**
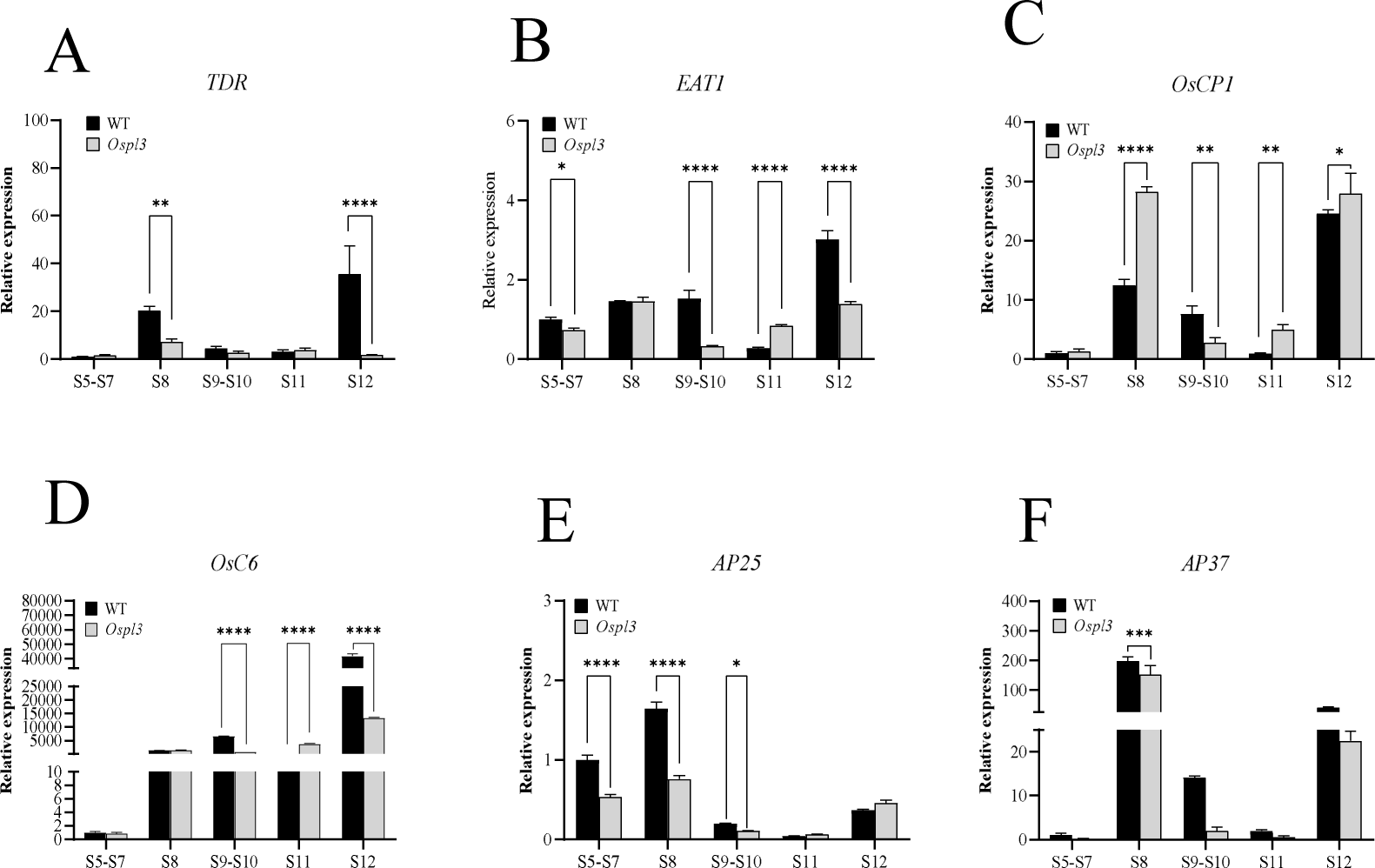
Expression analysis of the PCD marker genes from S5 to S12. Quantitative RT-PCR of *TDR* (A), *EAT1/DTD* (B), *OsCP1* (C), *OsC6* (D), *AP25* (E), and *AP37* (F).

### Expression Changes in Genes Involved in ROS Homeostasis in *Osplatz3* Anthers

In Arabidopsis, a highly sophisticated and redundant network of at least 289 genes is involved in regulating ROS levels (Gechev et al., 2006). Transcriptome profiles revealed that 229 annotated ROS-related genes in *Osplatz3* showed significant changes from S5 to S12 (Fig. 6). GO term analysis indicated a high enrichment of “oxidoreductase activity” and “monooxygenase activity” at different stages of anthers development, suggesting that *OsPLATZ3* may regulate ROS homeostasis (Supplemental Fig. S8). Only 2 genes were identified as responding to reactive oxygen species and oxidative stress, demonstrating significant alterations (Fig. 6, Supplemental Table S3). These data suggest that the *Osplatz3* severely inhibits ROS production, resulting in the expression of only very few ROS-responsive genes (Fig. 6).

**Figure 6.**
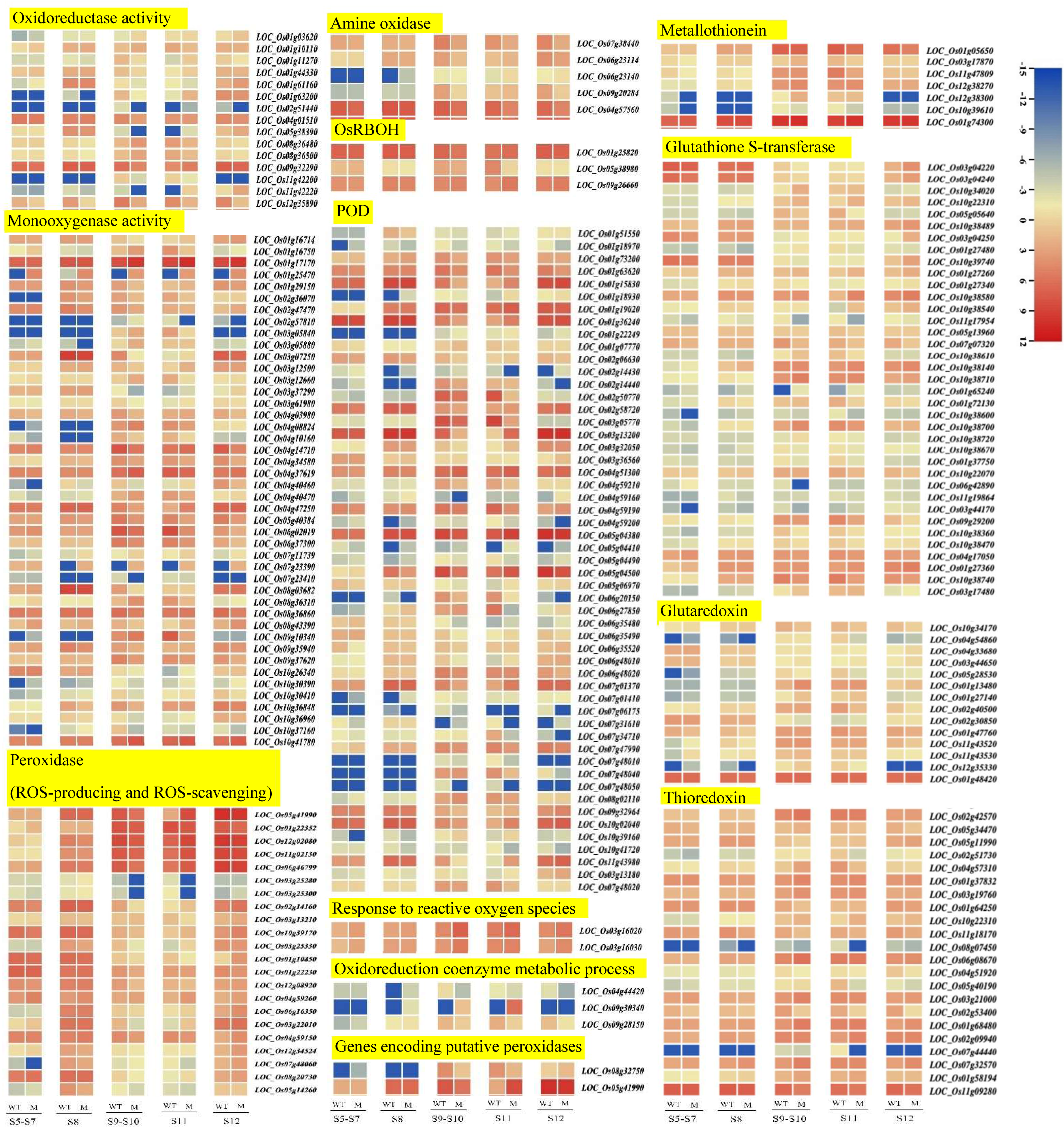
Heatmap of differential expression of 229 ROS-related genes in *Osplatz3*. Transcriptome profiles revealed that 229 annotated ROS-related genes in *Osplatz3* showed significant changes from S5 to S12, including 15 genes annotated for oxidoreductase activity, 44 genes annotated for monooxygenase activity, 2 genes annotated for response to reactive oxygen species and response to oxidative stress, 3 genes annotated for oxidoreduction coenzyme metabolic process, 7 metallothioneins, 37 glutathione S-transferases, 14 glutaredoxins, 22 thioredoxins, 2 genes encoding putative peroxidases, 53 POD genes, 22 peroxidase genes that cannot be distinguished between cell wall-bound peroxidases involved in ROS production and peroxidases required for ROS scavenging, as well as 5 amine oxidase genes, and 3 *OsBROHs*, which showed significant differential changes in their expression levels.

From S5 to S12 of anther development in *Osplatz3*, 7 metallothioneins; 37 glutathione S-transferases; 14 glutaredoxins; 22 thioredoxins; 2 genes encoding putative peroxidases, and 53 POD genes exhibited significant differential expression. Additionally, 22 peroxidase genes could not be distinguished between ROS production and scavenging, and their expression levels undergo significant differential changes (Fig. 6, Supplemental Table S3). These data suggest that the *OsPLATZ3* mutation affects the expression of ROS scavenging genes, and their protein products may be part of the ROS homeostasis network during anthers development.

The ROS levels in the *Osplatz3* anthers are consistently lower than those in the wild-type. Many changes in gene expression related to ROS production were observed in the transcriptome data of *Osplatz3*. In plants, ROS is mainly produced through the enzyme activity of NADPH oxidases bound to the plasma membrane, pH-dependent cell wall-bound peroxidases, and amine oxidases (Gechev et al., 2006). The expression levels of 3 *OsRBOH* genes and 5 amine oxidase genes in *Osplatz3* anther showed significant differences (Fig. 6, Supplementary Table 3). These data indicate that the mutation in *OsPLATZ3* has disrupted the ROS homeostasis in rice anther, highlighting its important role in regulating ROS homeostasis.

We corroborated the transcriptome data using qRT-PCR with independently generated RNA samples and focused on 8 differentially expressed genes (primers provided in Supplemental Table S1). The validated genes consist of metallothionein (*Os01g05650*, *Os12g38300*) (Supplemental Fig. S9, A-B), four peroxidases (*Os02g50770*, *Os05g04500*, *Os06g46799*, *Os12g02080*) (Supplemental Fig. S9, C-F), and two cytochromes (*Os06g07869* and *Os02g30100*) (Supplemental Fig. S9, G-H). Most of the proteins encoded by these genes are known or predicted to be involved in ROS homeostasis. The consistency between the changes observed from the transcriptome data analysis and qRT-PCR validates the reliability of the transcriptome results.

### OsPLATZ3 is associated with the promoter of ROS-Scavenging gene *OsAPX9*

To identify the downstream targets of OsPLATZ3, genes associated with ROS and exhibiting significant differences were chosen for qRT-PCR and promoter analysis. According to the results, *Os04g51300* is likely a downstream gene of OsPLATZ3. Based on gene annotation, *Os04g51300* is a plant ascorbate peroxidase domain-containing protein, and although the rice APX family has 8 members (Teixeira et al. 2006), *Os04g51300* is not among them. Additionally, *Os04g51300*, named as *OsAPX9* in our study, shows homology with Arabidopsis *APX4,* which possesses the capability to scavenge ROS. The qRT-PCR results showed significant upregulation of *OsAPX9* in *Osplatz3* from S9 to S12 compared to the wild-type (Fig. 7A). Promoter analysis of the *OsAPX9* gene revealed the presence of an AT-rich motif from –657bp to –628bp upstream of the start site (Fig. 7C).

**Figure 7.**
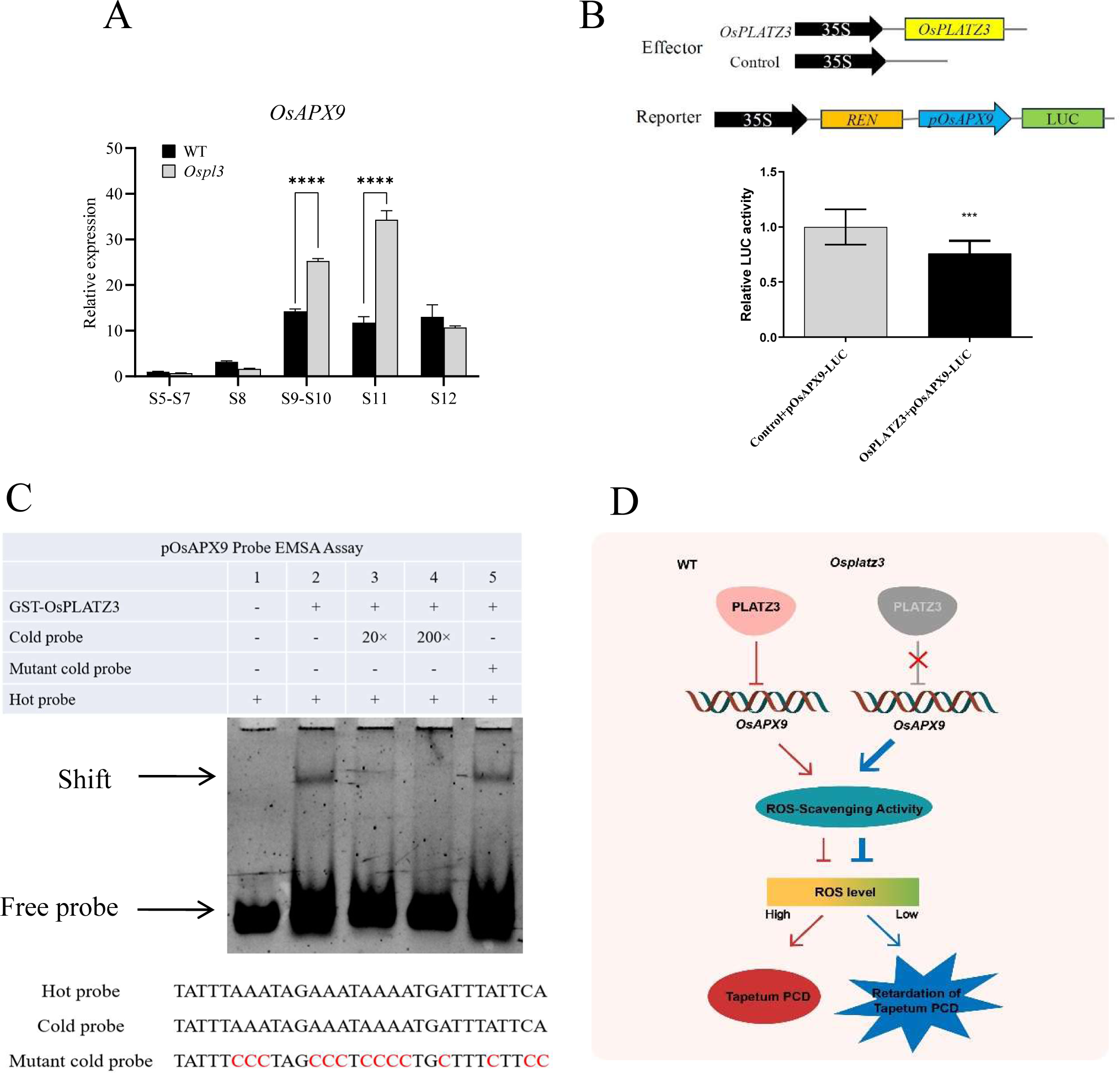
Regulation of downstream gene OsAPX9 by OsPLATZ3. A, qRT-PCR analysis of downstream gene OsAPX9 regulated by OsPLATZ3 in wild-type and *Osplatz3*. B, Dual-Luciferase Assay. C, EMSA Assay. D, Model of OsPLATZ3 regulating ROS levels through *OsAPX9* influencing the tapetum PCD.

The dual-luciferase assay was employed to investigate whether OsPLATZ3 had the ability to directly regulate *OsAPX9*. The results demonstrated a significant decrease in the luciferase activity of 62SK-OsPLATZ3 and *pOsAPX9*-LUC compared to the control group. The findings indicate that OsPLATZ3 may have specific binding sites with *OsAPX9* promoter, and OsPLATZ3 can targetly inhibit the expression of *OsAPX9*, suggesting that *OsAPX9* may be a downstream target gene of OsPLATZ3 (Fig. 7B).

The Electrophoretic Mobility Shift Assay (EMSA) was conducted to investigate the direct binding of OsPLATZ3 to the AT-rich motif within the *OsAPX9* promoter. The results showed that the GST-OsPLATZ3 recombinant protein directly interacts with the AT-rich motif within the *OsAPX9* promoter sequence. Competitive inhibition assays using excess unlabeled probe DNA revealed weakened binding of OsPLATZ3 to the probe, indicating specificity of the interaction. Nucleotide site mutation experiments further confirmed the specificity of the binding, demonstrating that OsPLATZ3 failed to bind to the mutant probe (Fig. 7C). These results confirm that *OsAPX9* is a target gene regulated by OsPLATZ3, which may result in increased activity of ROS scavenging enzymes and decreased ROS levels in *Osplatz3* compared to the wild-type (Fig. 7D).

## DISCUSSION

### OsPLATZ3 is a newly identified Tapetal PCD regulatory factor

The PLATZ transcription factor family plays an indispensable role in the growth and development of plants and their responses to abiotic stress. Several PLATZ family genes have been studied in Arabidopsis, maize, wheat and rice, with functions related to seed size (Wang et al., 2019; Zhou and Xue, 2019), endosperm formation (Li et al., 2017; Li et al., 2021), ROS regulation (Yamada et al., 2019), responses to high salt and drought stress (Liu et al., 2020; Liu et al., 2023), and hormone signaling pathways (Timilsina et al., 2022; Liu et al., 2023; Zhang et al., 2023). In this study, the rice *OsPLATZ3*, belonging to the PLATZ gene family, is predominantly expressed in the tapetum during late anther development. The delayed tapetal PCD observed in *Osplatz3* led to male sterility in rice. Unlike other PLATZ family members, *OsPLATZ3* primarily regulates tapetum PCD in rice.

After meiosis, the tapetum of rice undergoes PCD, which is believed to provide nutrients and signaling molecules to support pollen development (Li et al., 2006; Li et al., 2011). Timely tapetal PCD is crucial for normal pollen development and the formation of pollen walls. The tapetum PCD is regulated by multiple transcription factors in rice, and several conserved transcription factors related to tapetal PCD have been identified, including GAMYB, UDT1, TIP2, TDR, EAT1/DTD, PTC1, OsMADS3, as well as their downstream *DTC1* and effectors *OsC6*, *OsCP1*, *OsAP37*, and *OsAP25* (Tariq et al., 2022). These findings depict a complex transcriptional and post-transcriptional regulation network centered around TDR and EAT1, which is crucial for the precise occurrence of tapetum PCD and male fertility.

Our study shows that *OsPLATZ3* plays an important role in the tapetum PCD by inhibiting the ROS level. Transcriptome data analysis and qRT-PCR revealed a notable downregulation of *TDR* and *EAT1* in the *Osplatz3*, along with significantly different expression of *OsCP1* and *OsC6*, which both regluated by *TDR* as well (Fig. 5). *OsCP1* plays a crucial role in intracellular protein degradation and serves as a marker of PCD. Mutation leading to loss of *OsCP1* function result in the collapse of microspores after tetrad release (Lee et al., 2004). *OsC6* is a plant lipid transfer protein primarily expressed in the tapetal cells, and its silencing results in reduced pollen fertility (Zhang et al., 2010). This indicates that the mutation of *OsPLATZ3* impacts the expression of *TDR*, *EAT1*, *OsCP1*, and *OsC6*, leading to delayed tapetum degeneration and affecting pollen fertility. We speculate that they may be located downstream of OsPLATZ3 or interact with OsPLATZ3. Furthermore, OsPLATZ3 also suppresses the expression of AP25 and AP37 (Fig. 5), which have the ability to trigger cell death when ectopically expressed. Thus, we theorize that the absence of these cell death-inducing proteins in *Osplatz3* leads to delayed tapetum degeneration. In conclusion, we speculate that OsPLATZ3 may have a distinct role in the regulatory network of tapetum degeneration through the central regulatory pathway involving TDR and EAT1. Currently, there is insufficient research on the tapetum degeneration regulatory network of OsPLATZ3, and further studies are required to determine the specific role of OsPLATZ3 in this network.

### *OsPLATZ3* regulates ROS homeostasis during the late anther development

ROS has emerged as a crucial indicator for the activation of plant PCD and is involved in various instances of plant cell death (Hoeberichts and Woltering, 2002). The tapetum degeneration is believed to be triggered by the PCD during the late stages of anther development (Varnier et al., 2005). Some ROS molecules, such as superoxide radical and hydrogen peroxide, are key regulators of plant cell death (Hu et al., 2011). For example, OsMT-I-4b is highly induced by OsMADS3, maintaining ROS homeostasis and triggering tapetum PCD (Hu et al., 2011). The DTC1 protein interacts with the ROS scavenger OsMT2b, inhibiting ROS scavenging activity, and becomes a key regulatory factor in tapetum PCD (Yi et al., 2016).

During the development of anther in wild-type rice, the tapetum PCD can be first detected at S8 and a strong PCD signal appears at S9, indicating the timely production of ROS, especially superoxide anion, is related to the initiation of tapetum PCD (Li et al., 2006; Zhang and Wilson, 2009). In *Osplatz3*, we found that tapetum PCD is delayed and disordered, and the tapetum PCD signal continues from S8 to S11, while in the wild-type, the tapetum PCD signal terminates at S10 (Fig. 4A). Consistently, in *Osplatz3*, the ROS levels are lower from S8 to S12 compared to the wild-type (Fig. 4, B and C). In addition, the expression levels of *OsAPX9* are significantly upregulated from S9 to S12 (Fig. 7A), indicating enhanced OsAPX9 enzyme activity and improved ROS scavenging capacity, leading to lower ROS levels in *Osplatz3* compared to the wild-type (Fig. 4, B and C). The decrease in ROS levels in the *Osplatz3* anthers causes delayed and disordered tapetum PCD, resulting in male sterility, demonstrating the crucial role of maintaining ROS homeostasis in pollen development (Fig. 7D). *OsAPX9* is similar to Arabidopsis *APX4* (AT4G09010). *APX4* is situated in the chloroplast, and *apx4* exhibits leaf chlorosis, heightened H_2_O_2_ accumulation, and reduced soluble APX activity in seedlings. These results suggest that APX4 is involved in ROS scavenging processes in chloroplasts (Wang et al., 2014). It is hypothesized that *OsAPX9* may function similarly to *APX4*, given the shared phenotypes in *apx4*, such as leaf chlorosis and ROS scavenging, being observed in *Osplatz3* as well. However, further in *vivo* experiments are necessary to verify if the recombined OsAPX9 protein can support the biochemical activity of OsAPX9 in scavenging H_2_O_2_, and whether the knockout of *OsAPX9* leads to the restoration of pollen fertility and an increase in superoxide anion accumulation in transgenic lines.

In *Osplatz3*, we also observed a significant enrichment of genes associated with ROS-associated oxidoreductase activity and monooxygenase activity during the process of anther development. The expression levels of 229 genes with potential roles in the ROS homeostasis network showed significant changes at different stages of anther development in *Osplatz3* (Supplemental Fig. S7; Supplemental Table S3). This suggests that *OsPLATZ3* may play an important role in regulating ROS homeostasis during anthers development. Compared to the wild-type, *Osplatz3* showed lower ROS accumulation at various stages of anther development from meiosis onwards (Fig. 4, B and C). The reduction in ROS levels may be related to the increased expression of ROS-scavenging genes. In other words, the increase in the activity of OsAPX9 in *Osplatz3* may only partially contribute to the decrease in ROS levels in the later anther development.

In conclusion, we found that OsPLATZ3 exhibits relatively high expression levels at various stages of rice anther development in this study. Mutation of *OsPLATZ3* leads to lower ROS levels in the late stage of anther development compared to the wild-type, causing delayed tapetum PCD and male sterility, indicating that OsPLATZ3 is a novel member of the PLATZ family proteins in the tapetum PCD regulatory network. OsPLATZ3 regulates ROS homeostasis by suppressing the expression of ROS scavenging genes including OsAPX9, thereby controlling tapetum PCD in rice. Our study provides new insights into the function of the PLATZ family proteins and expands our understanding of the molecular mechanisms that control ROS homeostasis and tapetum PCD in rice.

## MATERIALS AND METHODS

### Identification of *Osplatz3* mutants and growth conditions

Seeds from rice lines 1C-04607 (*Osplatz3-1*) and 4A-02615 (*Osplatz3-2*) were germinated, and seedlings were grown for 10 d at 28–30°C in a greenhouse (16/8 h light–dark cycle). Each plant was grown in normal rice culture solution. DNA was extracted from individual plants for mutant identification using primers (Supplemental Table S1). After identification, potted plants were grown and managed using conventional methods. Representative panicles with uniform development were selected as experimental materials for each rice genotype.

### Total RNA extraction, complementary DNA (cDNA) synthesis, and qRT-PCR

Total RNA extraction, complementary DNA (cDNA) preparation, and qRT-PCR were carried out following the previous description for rice anther (Li et al., 2013). All gene-specific primer sequences used in this study are listed (Supplemental Table S1). The amplification of various genes was normalized to the expression of *OsActin1* and their relative expression levels were determined by the 2^−ΔΔCT^ method (Schmittgen and Livak, 2008). Each treatment was replicated three times.

### Histochemical GUS Assays and Microscopic Analyses

The GUS histochemical analysis was performed according to the previously described methods (Jefferson et al., 1987). For light microscopy analysis, tissues were fixed in a solution containing 50% ethanol, 5% acetic acid, and 3.7% formaldehyde, and then embedded in paraffin. Sample sections of 5 to 7 μm thickness were cut using a microtome (Leica) and observed under a microscope (OLYMPUS, BX51).

### In situ hybridisation analysis

Fresh inflorescences were fixed in 4% paraformaldehyde in 4°C overnight. Following dehydration in an ethanol series, the fixed tissues were transferred to 100% xylene and embedded in paraffin as described by Nguyen (Nguyen et al., 2010). Sections were collected on slides and prepared for pre-hybridization. Hybridization and probe detection were carried out following the method of Endo (Endo et al., 2009).

### Subcellular localization

The full-length CDS of OsPLATZ3 was cloned into the pCAMBIA1300-35S-N-GFP vector to generate the pCAMBIA1300-35S-OsPL1-GFP vector (primer is provided in Supplemental Table S1). After Agrobacterium-mediated transformation, tobacco leaves were infiltrated, and 48-76 hours later, the infiltrated leaves were excised and observed and photographed under a laser confocal scanning microscope (OLYMPUS, FV1000).

### TUNEL assays

Fresh anthers collected from different developmental stages (S7-S12) are fixed overnight in FAA (50% ethanol, 5.0% glacial acetic acid, 3.7% formaldehyde), dehydrated in graded ethanol, embedded in paraffin, and sectioned into 5μm thick slices. Anther transverse sections are stained with Safranin O (SO)/Fast Green and incubated at 42°C for 1-2 hours. TUNEL assay is performed using the DeadEnd Fluorometric TUNEL System (Promega). Following incubation, sections are counterstained with DAPI working solution (1 μg/mL) for 10 minutes and observed using a confocal microscope (LSM 800; ZEISS).

### NBT staining and ROS content determination

The tissue chemical detection of O^2•−^ generation rate in fresh rice anthers uses the nitro blue tetrazolium (NBT) staining method (Yi et al., 2016). The detection of total ROS levels in rice anthers uses the 2,7-dichlorofluorescein diacetate (DCFH-DA) fluorescence detection method (Zhao et al., 2018). Fresh rice anthers are treated with 10 μM DCFH-DA dissolved in 0.1% dimethyl sulfoxide at 30°C avoiding light for 30 minutes, then washed with HBSS (pH 7.2). The reaction of fluorescent probe (DCFH-DA) with ROS forms high-fluorescence DCF, and the fluorescence intensity is observed and imaged using a fluorescence microscope (OLYMPUS BX51). The fluorescence signal intensity is measured using ImageJ software. As technical replicates, five to six fresh anthers from different florets are detected for each treatment.

### Dual luciferase transcriptional activity assay

The full-length cDNA of OsPLATZ3, OsPLATZ3N, and OsPLATZ3C was cloned into the yeast GAL4 binding domain vector (pGreenII 62-SK-GAL4BD) according to the Dual-Luciferase Reporter Assay system (Hellens et al., 2005). The full-length cDNA of OsPLATZ3 was also cloned into the yeast GAL4/VP16 binding domain vector (pGreenII 62-SK-GAL4BD-VP16) as the effector. The pGreenII 0800-5×UAS-LUC served as the reporter vector. Additionally, the full-length cDNA of OsPLATZ3 was cloned into the downstream of the 35S promoter in the pGreenII 62-SK vector to construct the 62SK-OsPLATZ3 effector vector. The promoter of OsAPX9 was cloned upstream of the LUC reporter in pGreenII 0800-LUC to construct the pGreenII 0800-pOsAPX9-LUC reporter vector. Agrobacterium tumefaciens was separately transformed and the tobacco leaves were co-infiltrated in a specific ratio. The values of LUC and REN were measured and the expression ratio of firefly luciferase to the internal control Renilla luciferase was calculated.

### RNA sequencing analysis

The *Osplatz3* and *Nipponbare* wild-type at different development stages were used for transcriptome analysis based on auricle distance (AD) under natural growth conditions; AD is the distance between the auricles of the flag leaf (last leaf) and the second-to-last leaf, used as a non-destructive measurement to estimate the developmental stages of microspores. Representative panicles with uniform development were chosen as experimental materials for each rice genotype. Plant total RNA was extracted using the QIAprep Spin Miniprep Kit (Qiagen 27104). Three biological replicate RNA samples were independently prepared for each genotype. A total of 10 libraries from S5-S7, S8, S9-S10, S11 and S12 of the wild-type and *Osplazt3* anthers were constructed for the analysis of the transcriptome. All libraries were sequenced separately on the Shanghai Sangon Biotech. In short, clean sequencing reads were mapped to the rice reference genome (the Rice Genome Annotation Project (RGAP, Release 7)). Differential expression genes (DEGs; |fold change| ≥ 2 and adjusted p-value <0.05) were selected using DESeq2 software (Love et al., 2014). Gene Ontology (GO) enrichment analysis was conducted using GOseq.

### EMSA experiments

The full-length coding region of the OsPLATZ3 sequence was cloned into the pGEX-4T-1 vector (primers see Supplemental Table S1). The vector was then transformed into BL21(DE3) and Rosetta (DE3). Protein was purified using GST purification medium. The 6-FAM (FITC) labeled reagent kit was used to synthesize the hot probe, while the cold probe was generated from the unlabelled dimeric oligonucleotide. A double-stranded DNA probe containing the CCTTTTGGGG box of P-OsAPX9 was prepared using the pOsAPX9-FF and pOsAPX9-R primers (probe see Supplemental Table S1). The steps of the DNA binding reaction were performed following the previous method (Wang et al., 2002).

### Construct for CRISPR/Cas9 and transformation

CRISPR-P (http://cbi.hzau.edu.cn/crispr/) was utilized for the selection of specific single guide RNA (sgRNA) that targeted OsPLATZ3. The vector pYLCRISPR/Cas9 was employed, and two 23 bp fragments from the 1st and 2nd exon of OsPLATZ3 were inserted into the vector. Subsequently, *A. tumefaciens*-mediated transformation was applied to rice *calli* induced of the wild-type. Transgenic plants were identified through PCR amplification and sequence analysis.

### Accession Numbers

Accession Numbers Sequence data from this article can be found in the Rice Genome Annotation Project (RGAP) or GenBank/EMBL databases under the following accession numbers: *CP1* (*Os04g57490*), *TDR* (*Os02g02820*), *OsEAT1 (Os04g51070), AP25(Os03g08790), OsAP37(Os04g37570), Os*C6 (Os11g37280*), OsMT-I-4b(Os12g38300), OsAPX9 (Os04g51300)*

## Acknowledgments

This paper is dedicated to Prof. Ping Wu, despite his passing. The author sincerely thanks the anonymous reviewers for their comments on the manuscript. Special thanks to the Salk Institute Genomic Analysis Laboratory for providing the T-DNA mutant materials. The guidance of Prof. Fengyi Hu, Prof. Yi Zhang and Prof. Chunzhao Zhao is greatly appreciated.

## Author Contributions

Y.L. conceived and designed the experiments; J.W., F.T., L.L., X.C., X.S., P.X., S.Y., Y.L, Y.W., M.T., and L.Z. performed the experiments; Y.L., L.L. and X.C. analyzed the RNA-seq data; Y.L. wrote the article with contribution of all the authors.

## Supplemental Data

The following materials are available in the online version of this article.

**Supplemental Figure S1.**
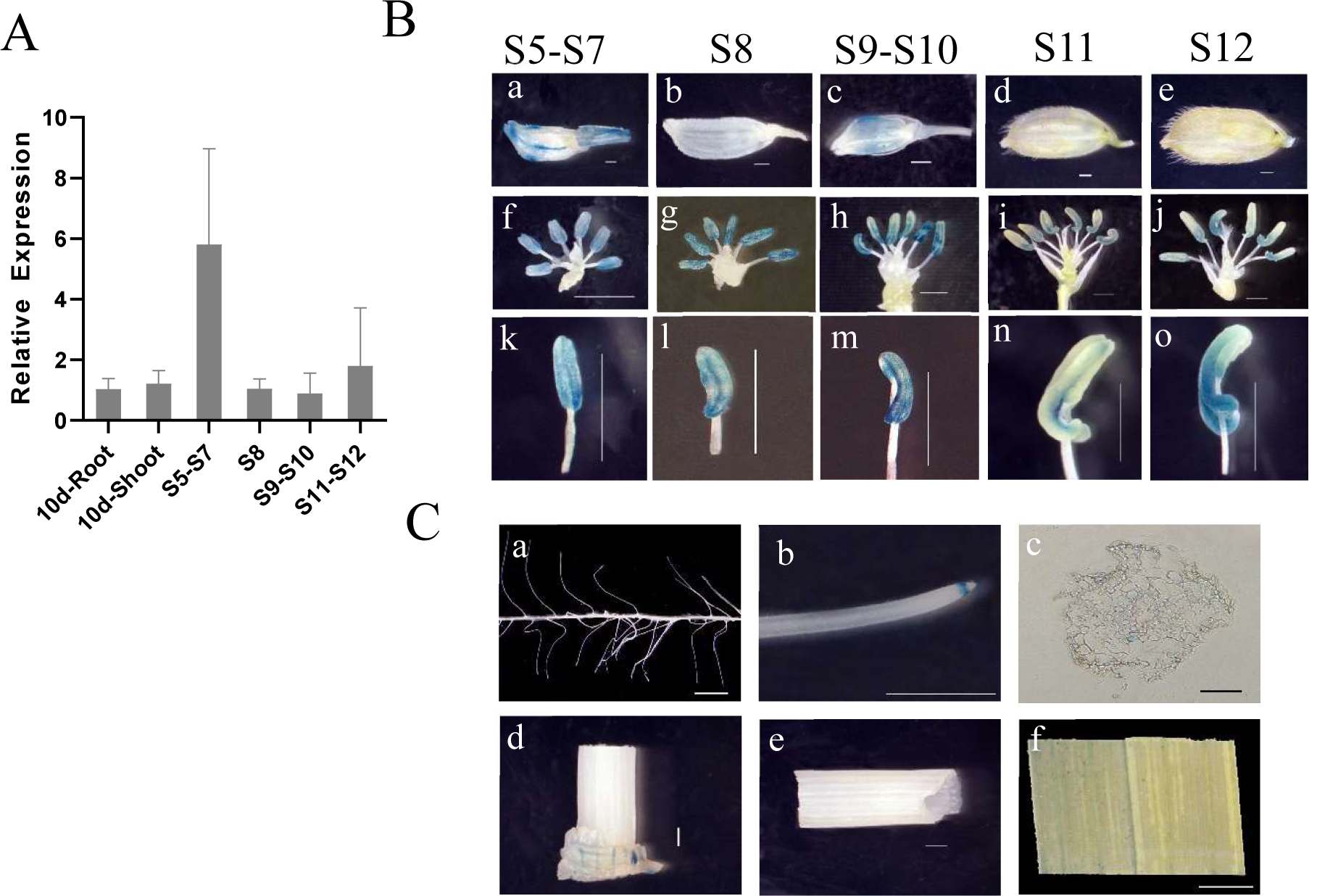
Expression analysis and GUS staining of *OsPLATZ3*. A, Expression analysis of *OsPLATZ3* in 10-day-old roots, 10-day-old shoot, and at anther developmental stages S5-S7, S8, S9-S10, S11, and S12. B, GUS activity analysis of OsPLATZ3 promoter in different developmental stages of rice inflorescence. S5-S7, MMC stage; S8, meiotic stage; S9-S10, uninucleate pollen stage; S11, binucleate pollen stage; S12, trinucleate pollen stage. C, GUS activity analysis of *OsPLATZ3* promoter in rice vegetative tissues. a, mature roots; b, root tip; c, transverse section of the root; d, root-shoot junction; e, stem; f, leaf. Bars = 1mm.

**Supplemental Figure S2.**
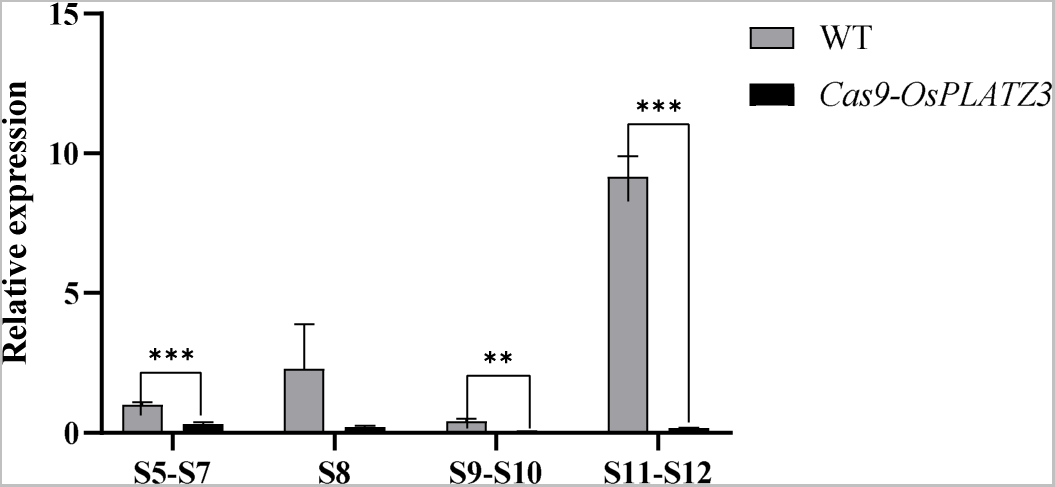
qRT-PCR analysis of *OsPLATZ3* expression levels during the S5-S12 of anther development in the gene editing material *Cas9-OsPLATZ3*.

**Supplemental Figure S3.**
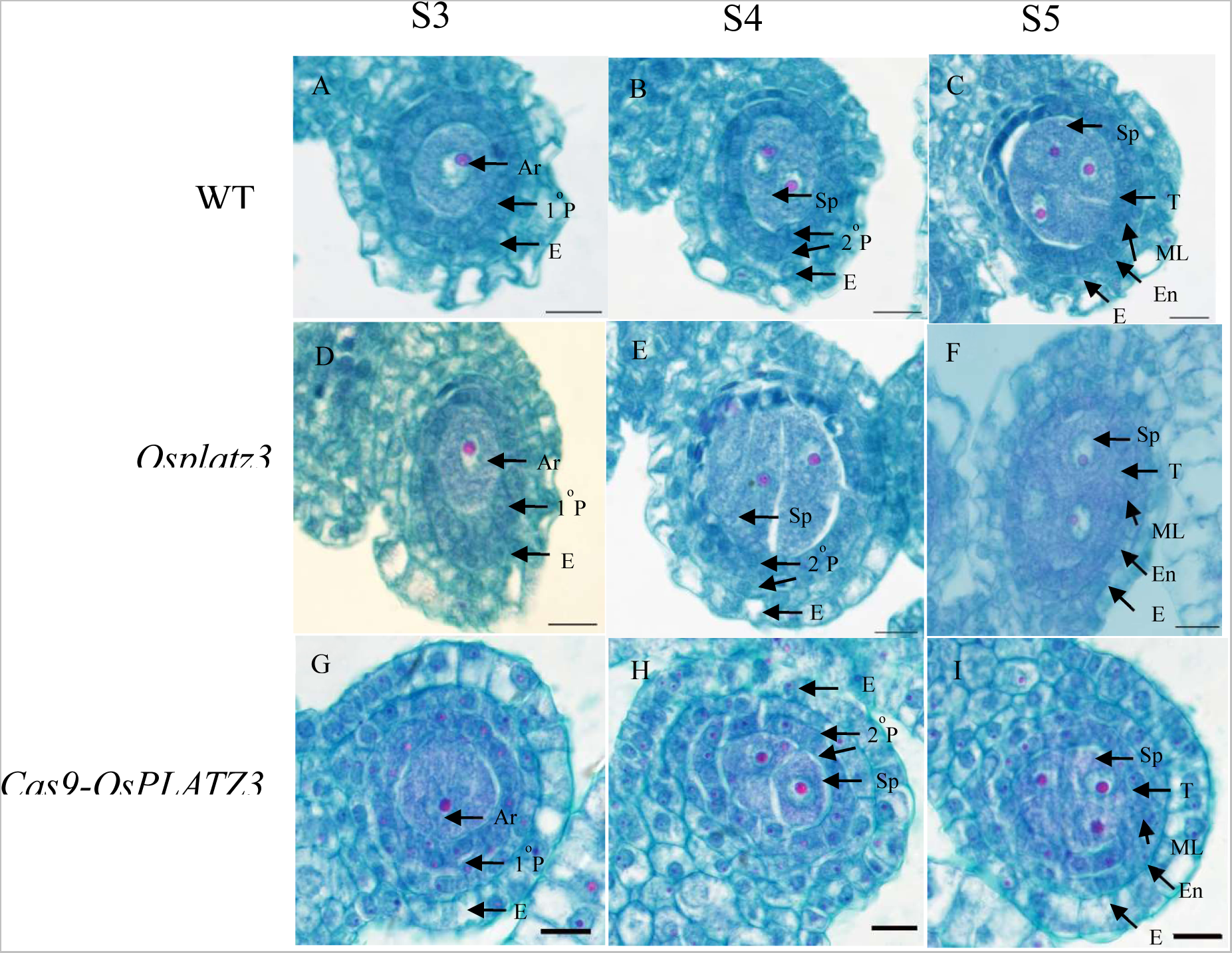
Transverse section comparison of wild-type, *Osplatz3*, and *Cas9-OsPLATZ3* during S3-S5 of anther development. Ar, archesporial cell; E, epidermis; 1°P, primary parietal layer; 2°P, secondary parietal cell layer; Sp, sporogenous cell; T, tapetum; En, endothecium; ML, middle layer. Bars = 10μm.

**Supplemental Figure S4.**
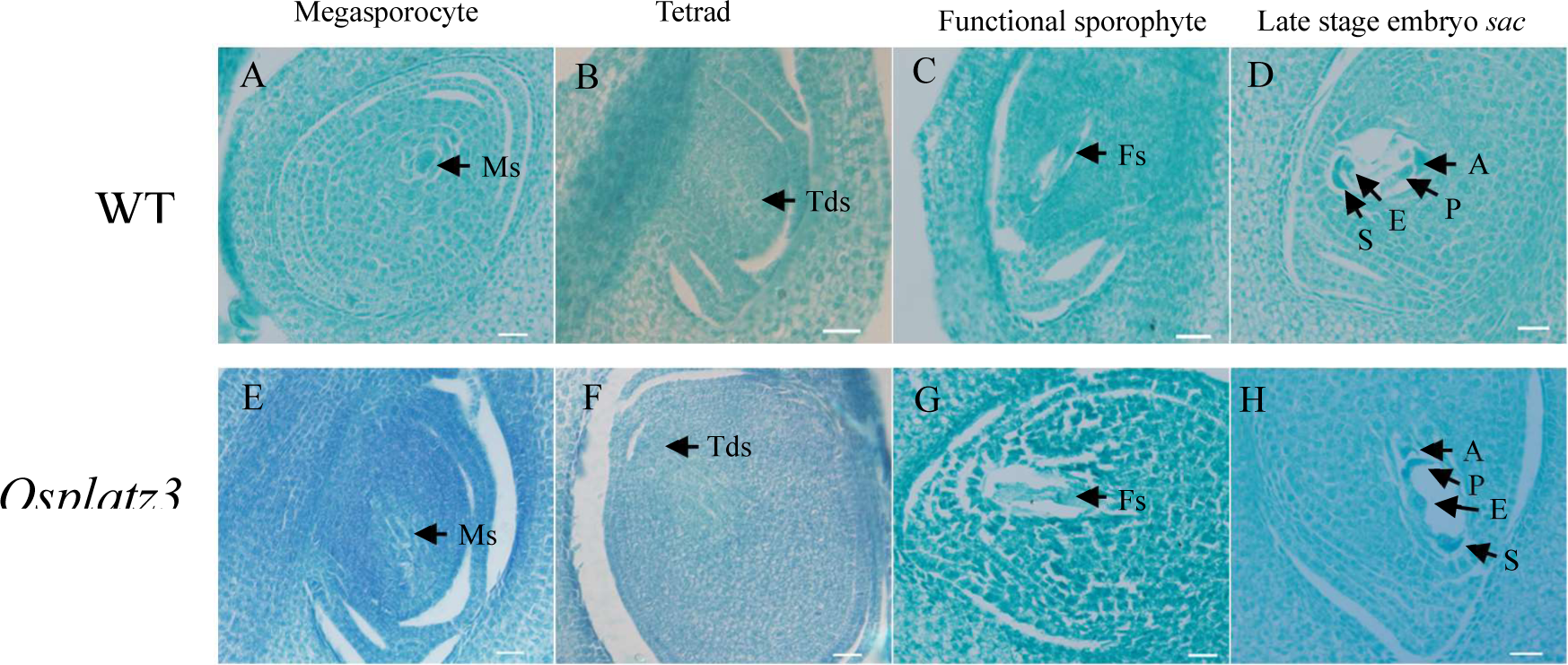
Transverse section comparison of wild-type and *Osplatz3* during the embryo *sacs* development. Ms, Megasporocyte; Tds, tetrads; Fs, Function sporophyte; A, Antipodal; P, Polar nuclei; E, Egg cell; S, Synergid. Bars = 10μm.

**Supplemental Figure S5.**
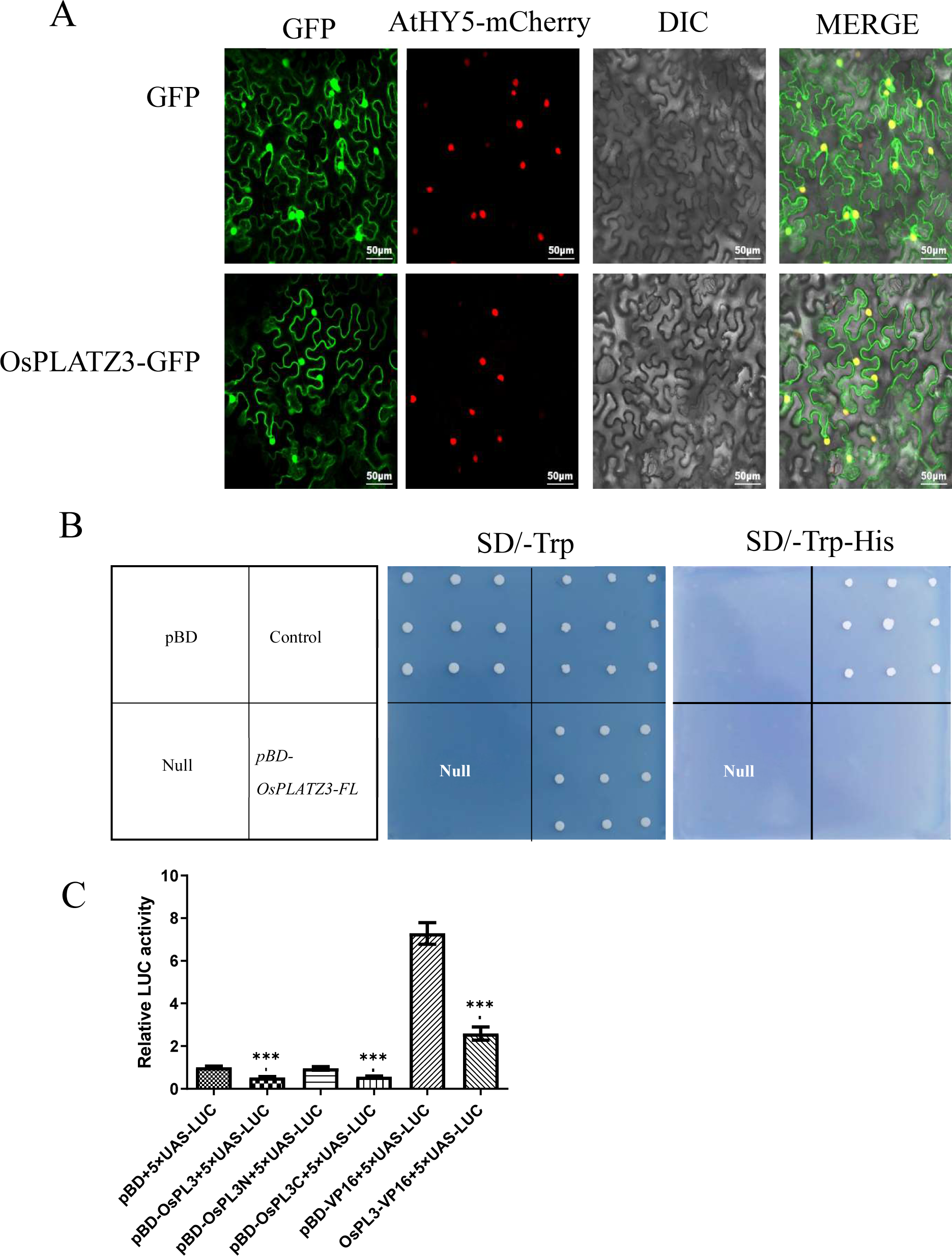
OsPLATZ3 is a nuclear-localized transcription repressor. A, Subcellular Localization of OsPLATZ3. Nucleus marker: pCAMBIA1300-35S-AtHY5-mCherry. Bars = 50 μm. B, Transactivation activity assay of OsPLATZ3 in yeast. pBD, negative control (pGBKT7 empty vector); Control, positive control; null, no sample. C, Transcriptional activation assay of OsPLATZ3. OsPLATZ3, abbreviated as OsPL3, and OsPLATZ3/VP16 fusion transcription factor significantly downregulated the activity of the reporter gene LUC, indicating that OsPLATZ3 is a transcriptional repressor.

**Supplemental Figure S6.**
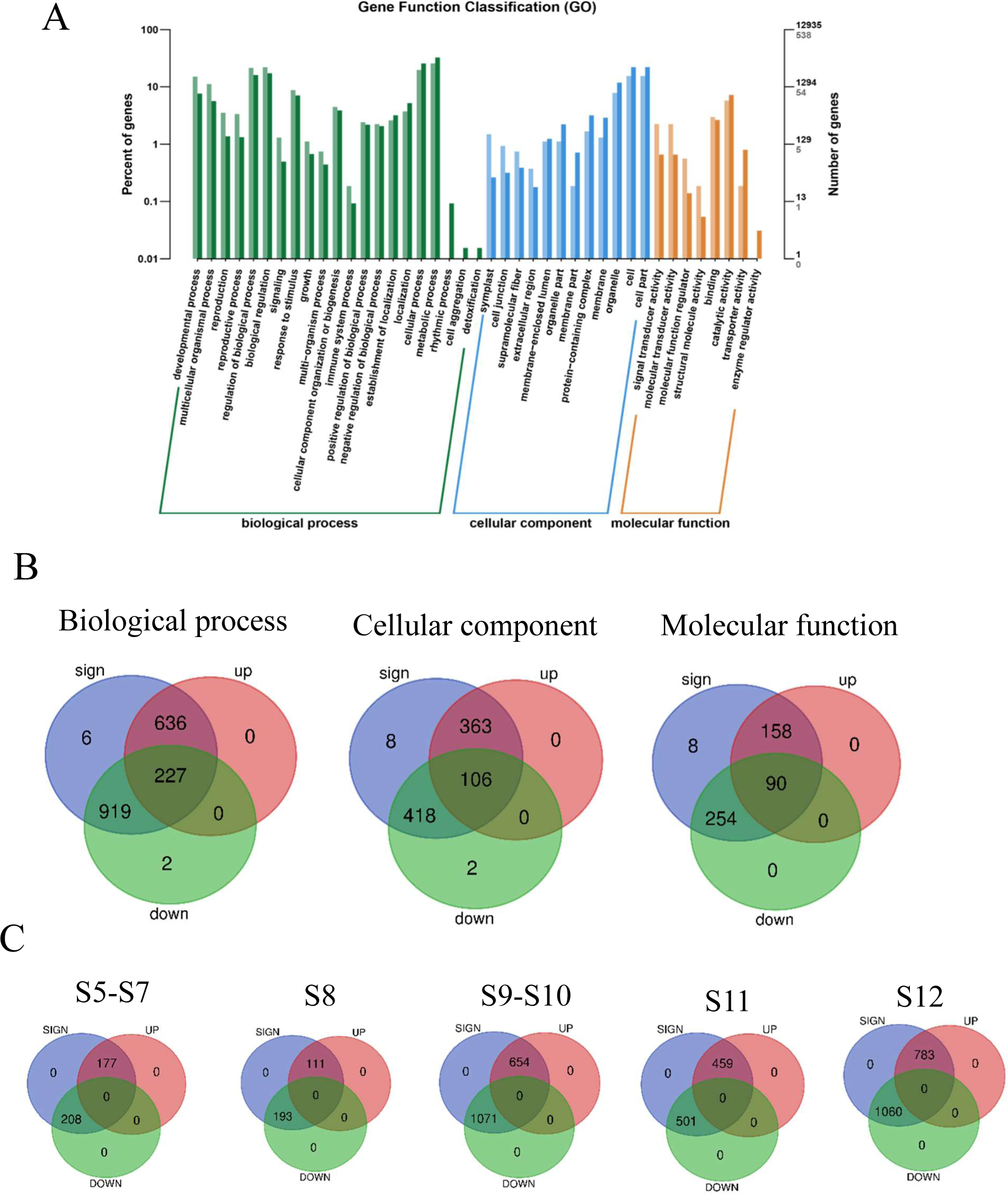
Transcriptome data analysis of *Osplatz3* at different stages of anther development. A, Classification of genes in transcriptome sequencing data. B, Three categories of genes showing significant changes in *Osplatz3*. C, Genes displaying significant differential expression in *Osplatz3* from S5 to S12 during anthers development.

**Supplemental Figure S7.**
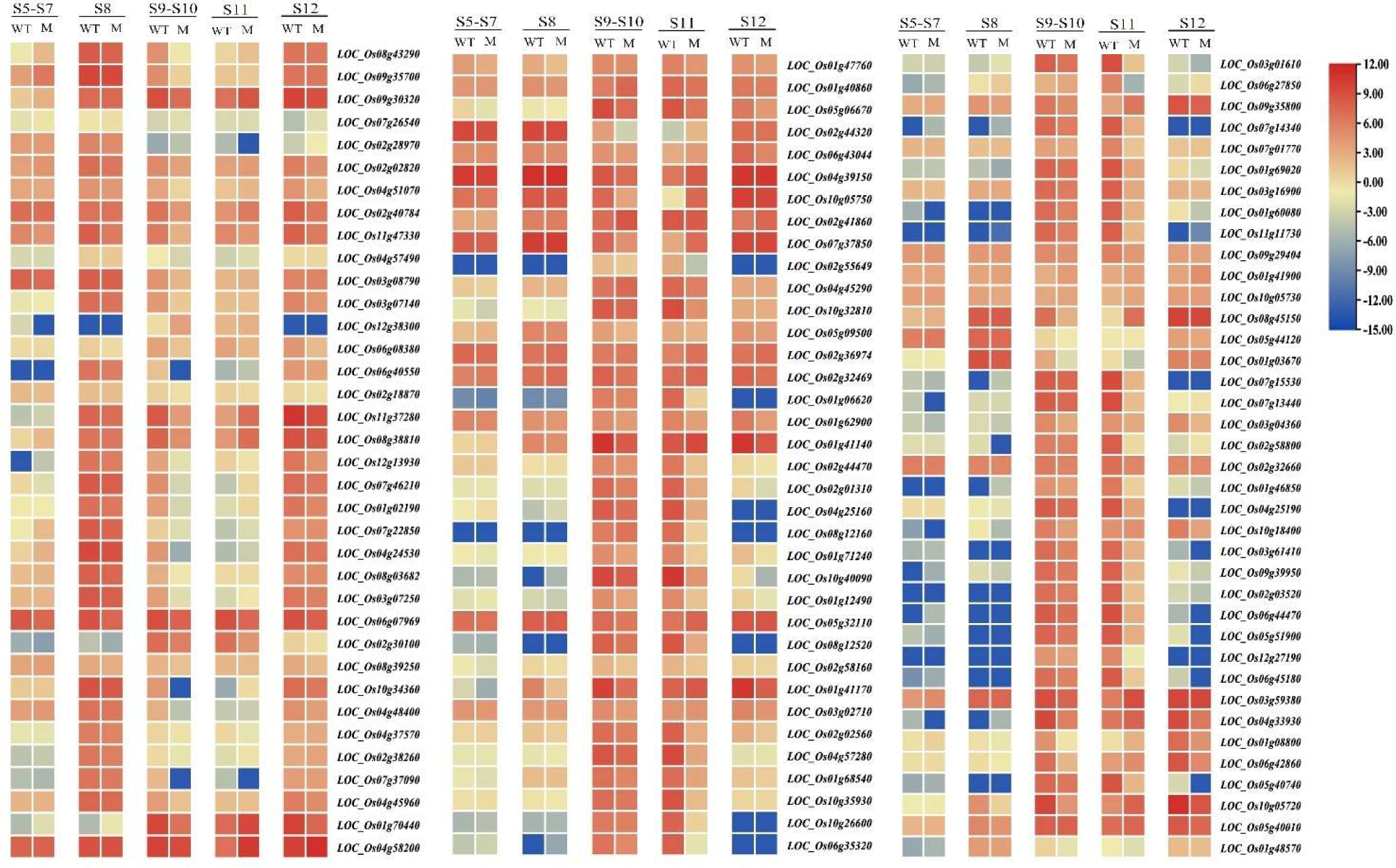
Heat map of pollen-specific genes with significant differential expression in *Osplatz3*. Transcriptome sequencing analysis revealed 112 genes associated with anthers development from S5 to S12 in *Osplatz3*, showing significantly different expression levels compared to the wild-type.

**Supplemental Figure S8.**
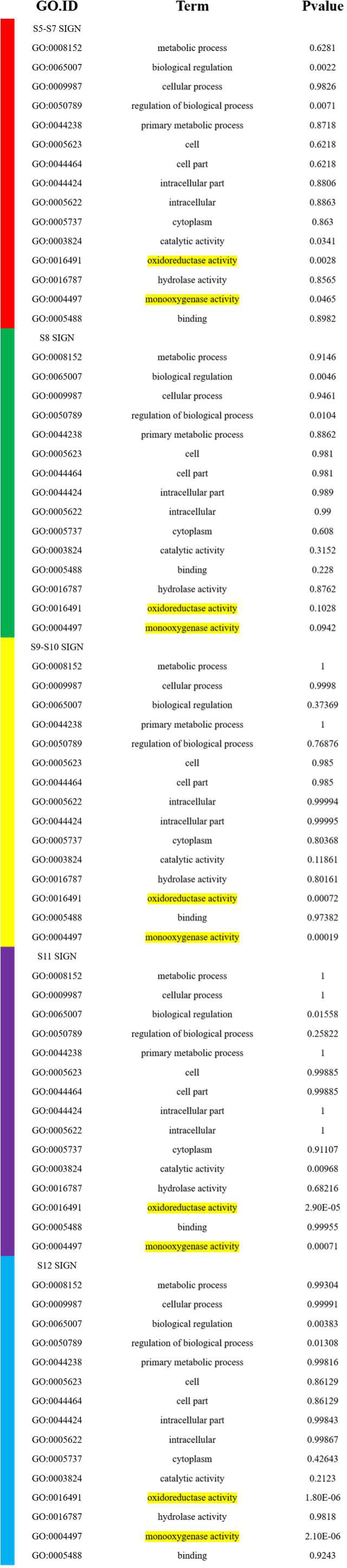
Enriched GO categories in the anther of *Osplatz3* from S5-S12. The gene ontology (GO) categories “Oxidoreductase activity” and “Monooxygenase activity” are highly significantly enriched (highlighted) in *Osplatz3*. The colors red, green, yellow, purple, and blue represent S5-S7, S8, S9-S10, S11, and S12 of rice anthers respectively.

**Supplemental Figure S9.**
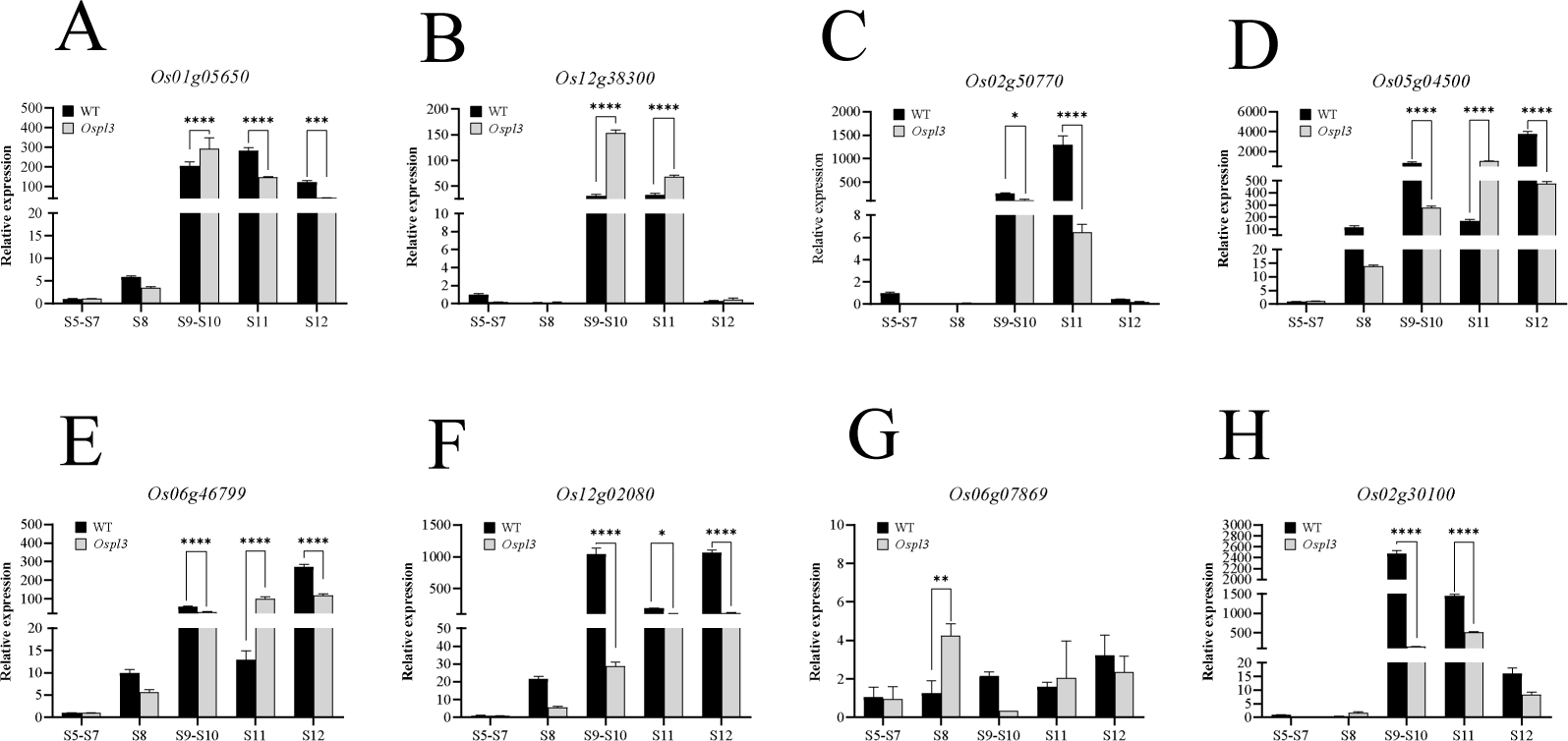
Expression analysis of ROS-related genes. Quantitative RT-PCR of two *MTs Os01g05650* (A), *Os12g38300* (*MT-I-4b)* (B), four peroxidase genes *Os02g50770* (C), *Os05g04500* (D), *Os06g46799* (E), *Os12g02080* (F), two cytochrome genes *Os06g07869* (G) and *Os02g30100* (H).

**Supplemental Table S1.**
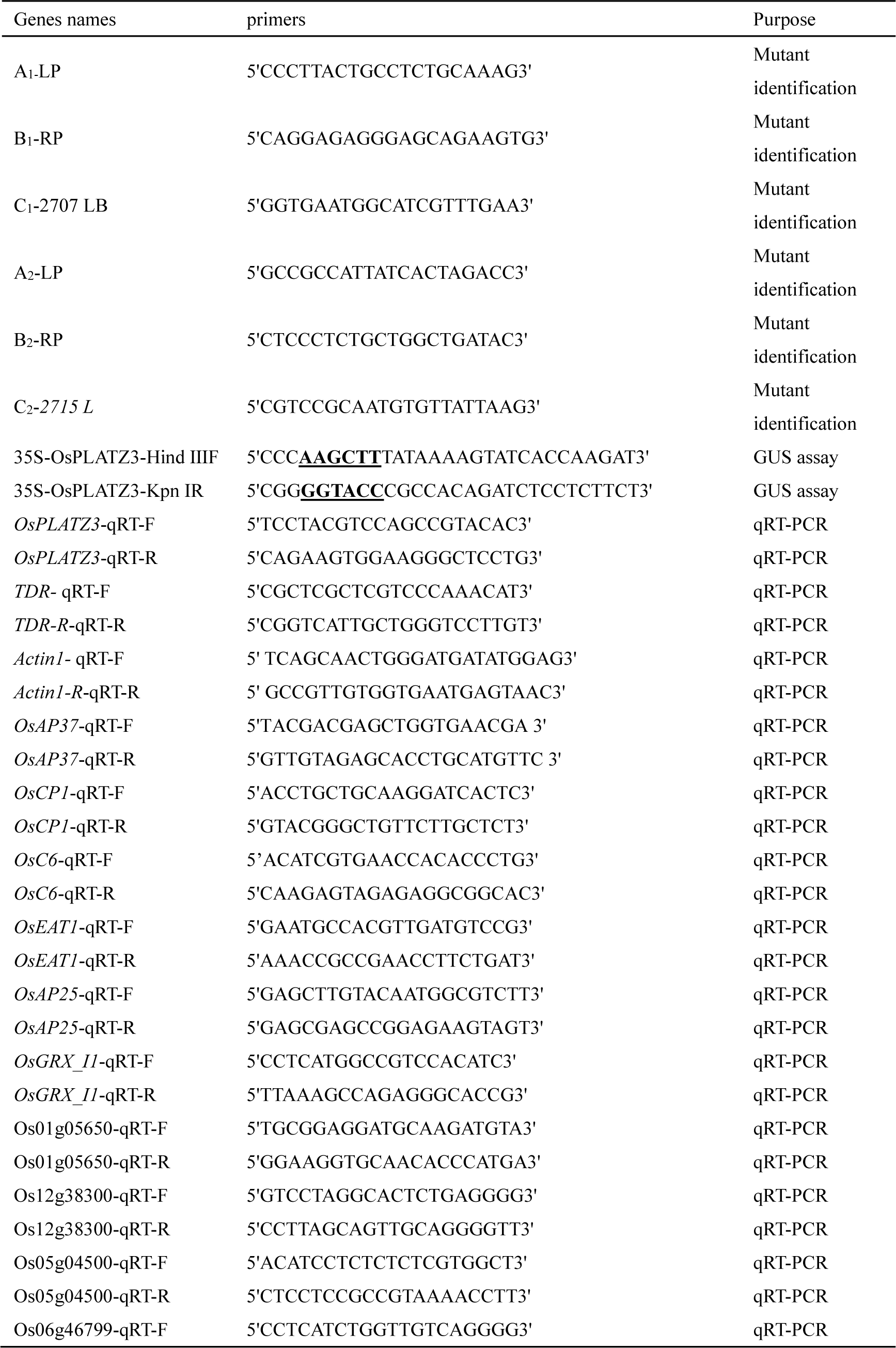
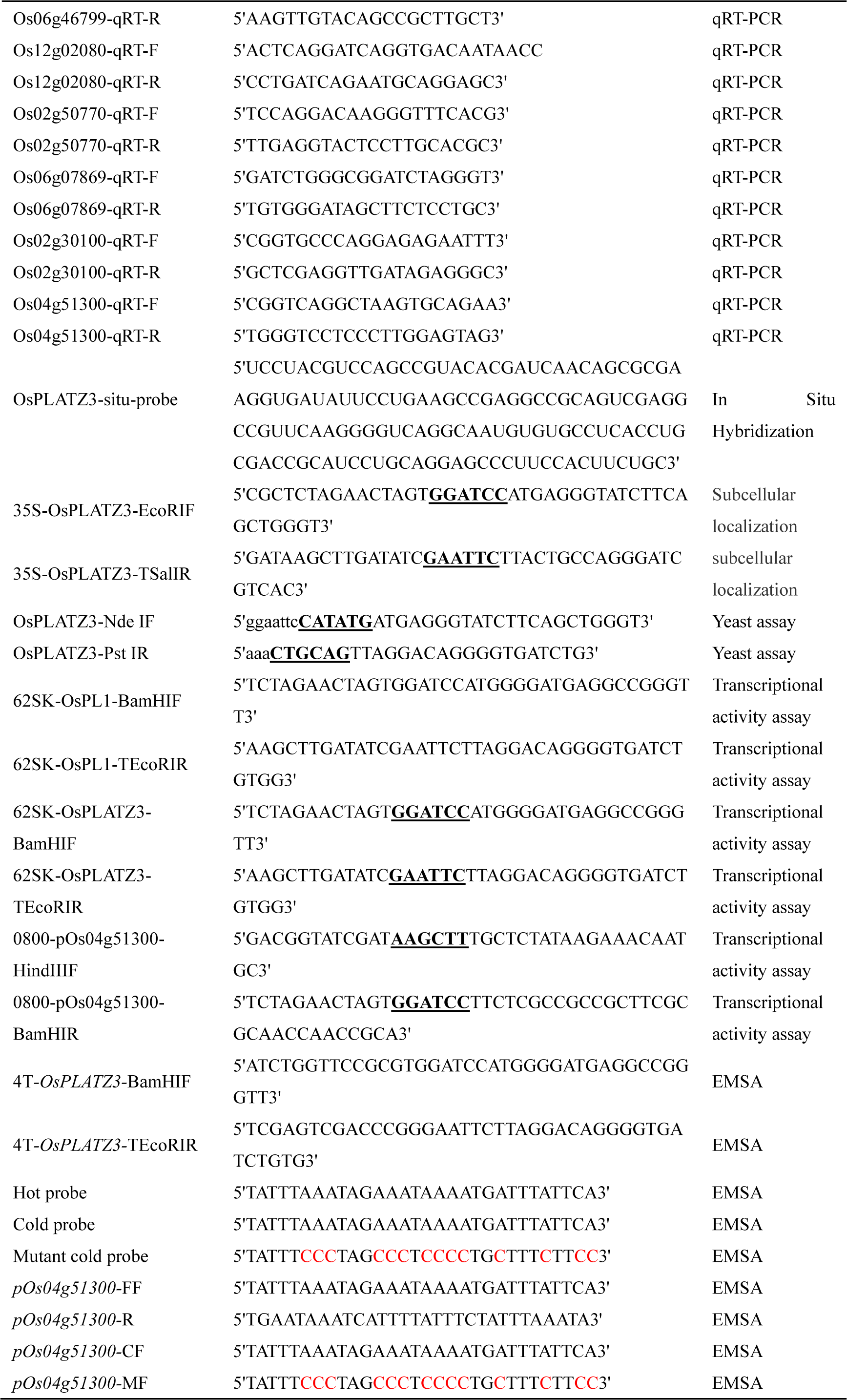
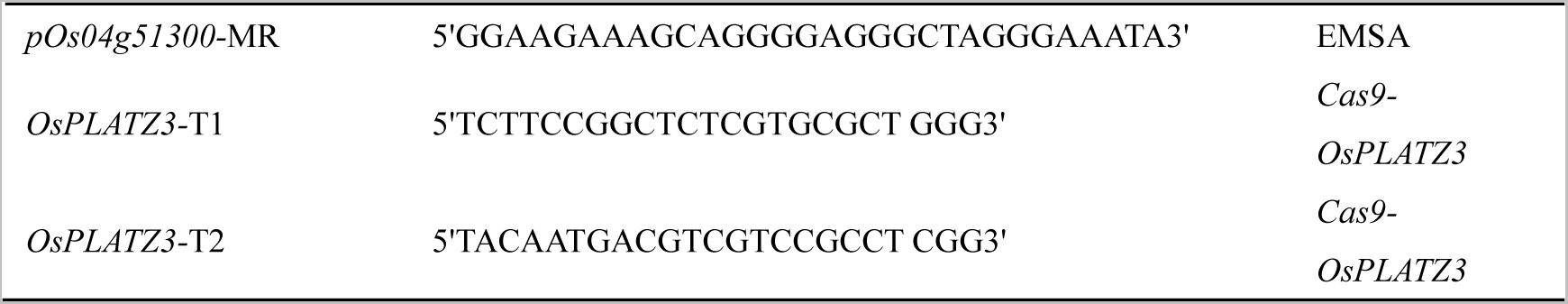
The relevant primers and probes used in the study.

**Supplemental Table S2.**
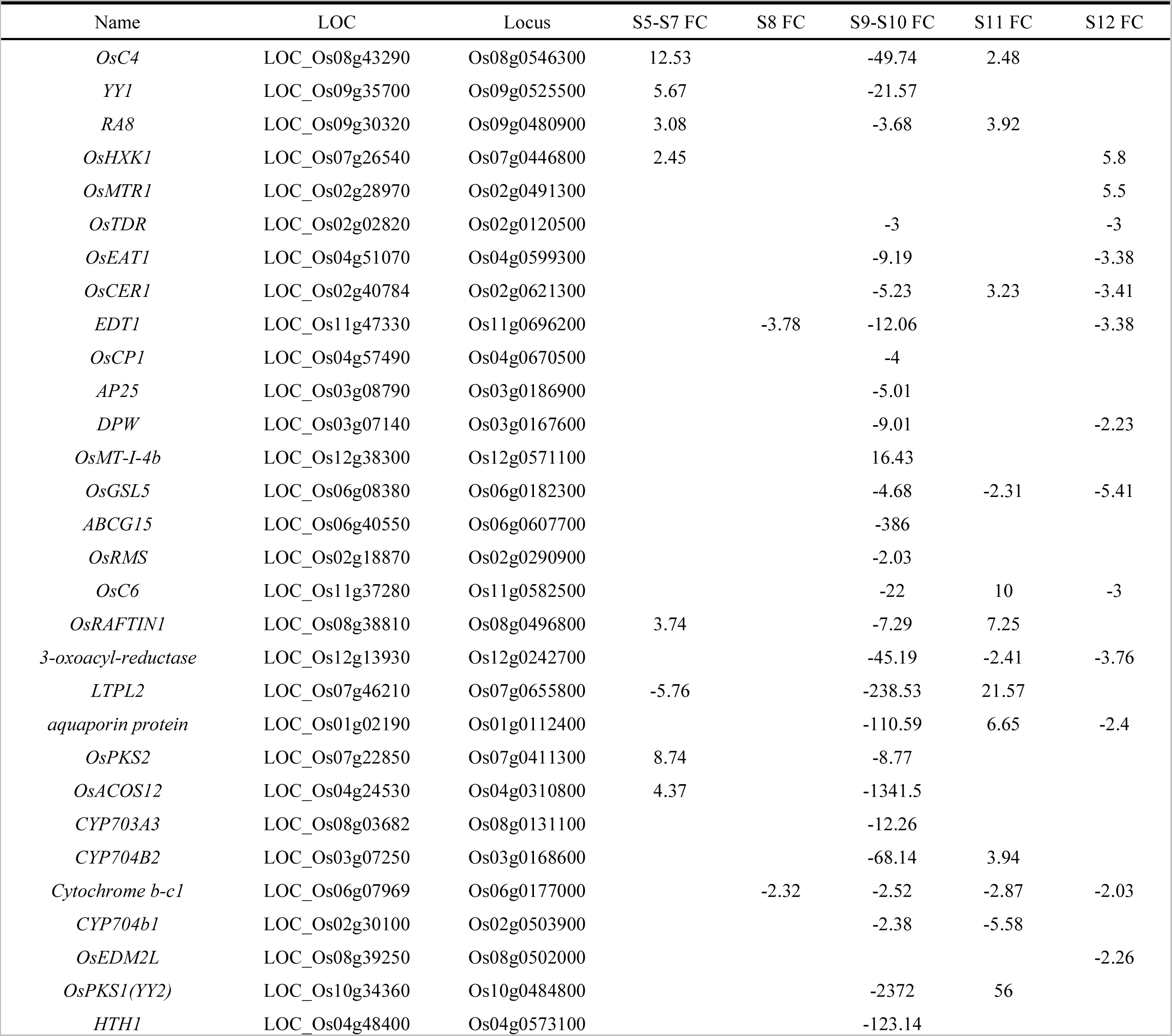
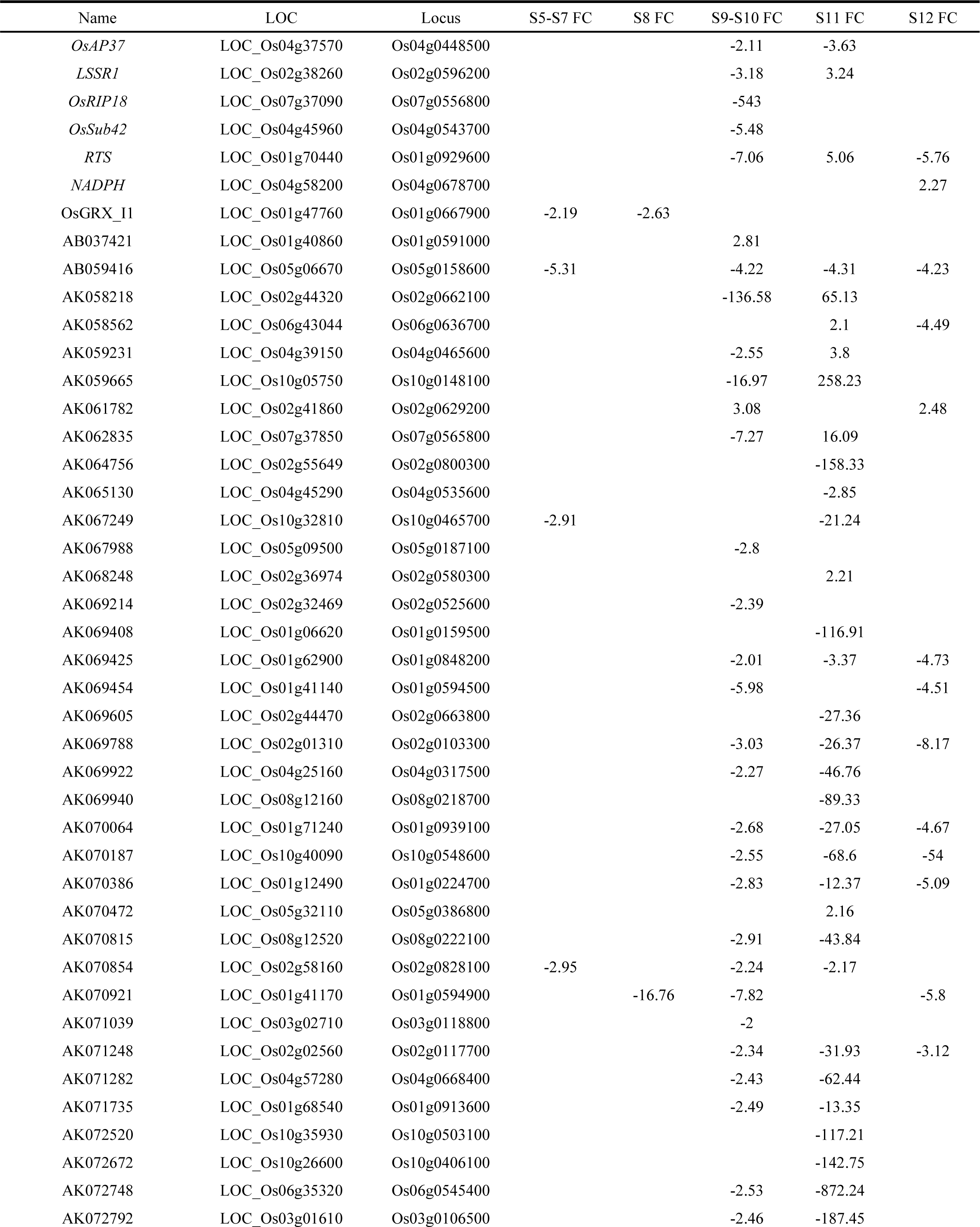
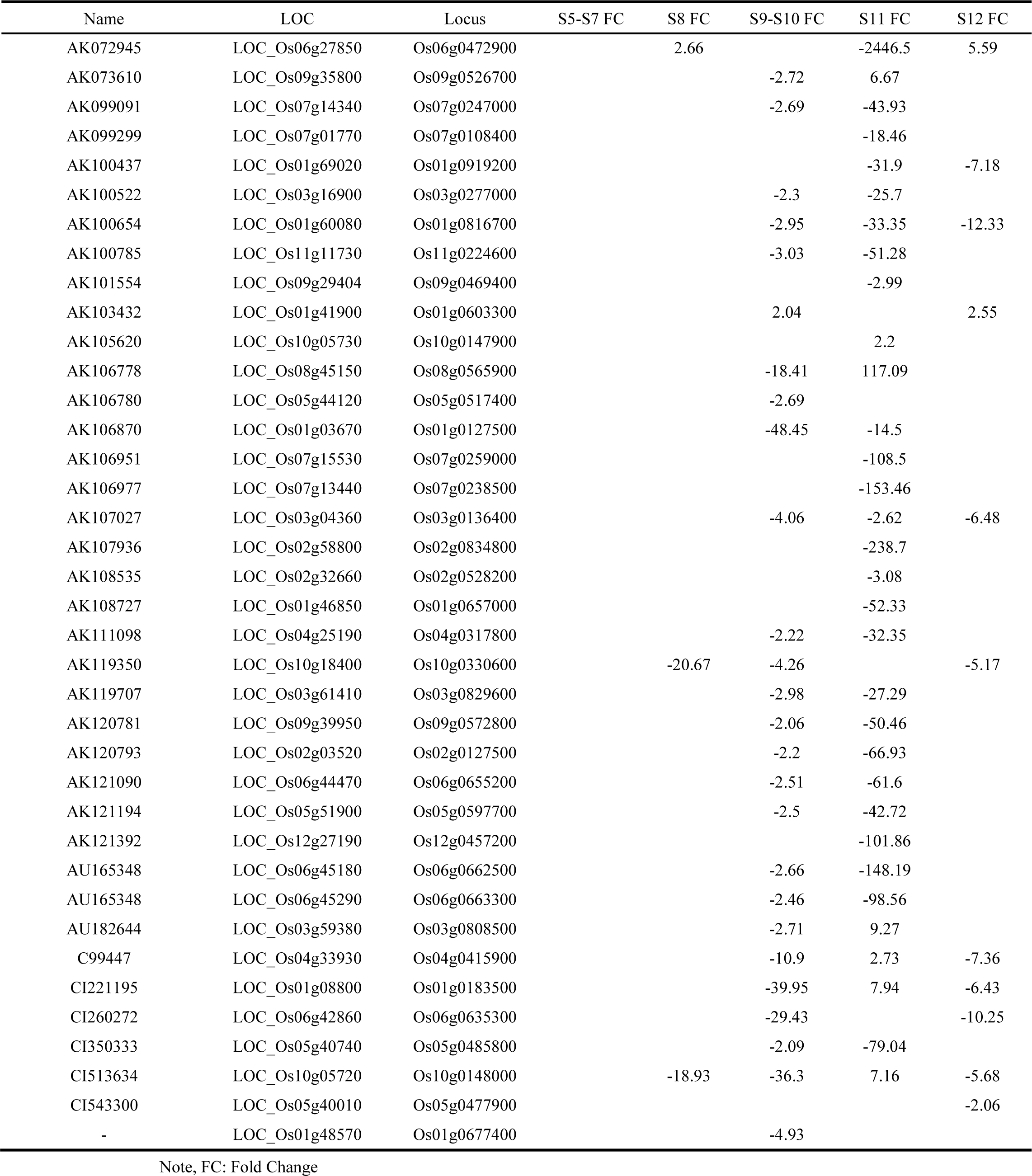
112 significantly differentially expressed anther-specific genes from S5 to S12 in *Osplatz3*.

**Supplemental Table S3.**
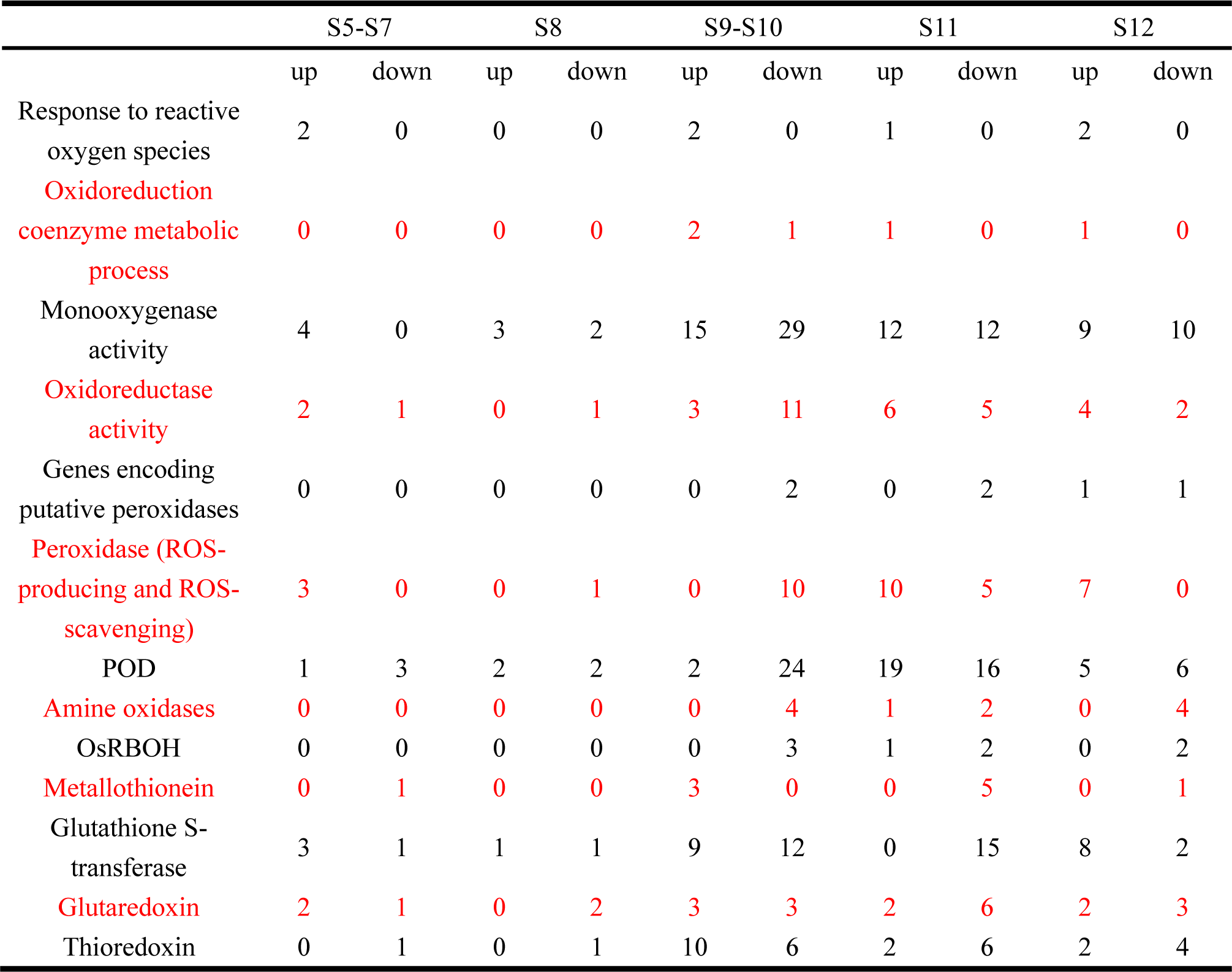
The total count of genes related to ROS homeostasis significantly upregulated and significantly downregulated has been statistically analyzed from S5 to S12 in *Osplatz3*.

